# Structural RNA components supervise the sequential DNA cleavage in R2 retrotransposon

**DOI:** 10.1101/2023.04.07.536001

**Authors:** Pujuan Deng, Shun-Qing Tan, Qi-Yu Yang, Han-Zhou Zhu, Lei Sun, Zhangbin Bao, Yi Lin, Qiangfeng Cliff Zhang, Jia Wang, Jun-Jie Gogo Liu

## Abstract

Retroelements are the widespread jumping elements considered as major drivers for genome evolution, which can also be repurposed as gene-editing tools. Here, we determined the cryo-EM structures of eukaryotic R2 retrotransposon with ribosomal DNA target and regulatory RNAs. Combined with biochemical and sequencing analysis, we revealed two essential DNA regions, Drr and Dcr, required for R2 recognition and cleavage. The association of 3’ regulatory RNA with R2 protein accelerates the first-strand cleavage, blocks the second-strand cleavage, and initiates the reverse transcription starting from the polyA tail. Removing 3’ regulatory RNA by reverse transcription allows the association of 5’ regulatory RNA and initiates the second-strand cleavage. Our work explained the DNA recognition and supervised sequential retrotransposition mechanisms by R2 machinery, providing novel insights into the retrotransposon and application reprogramming.

## Introduction

Retrotransposons are widespread in all three domains of life (Bonen and Vogel, 2001; Ferat and Michel, 1993; Kojima and Fujiwara, 2005; Lambowitz and Belfort, 2015; Xiong and Eickbush, 1988). Particularly in eukaryotes, retrotransposons have expanded via the copy-and-paste mechanism to shape both genes and the genomic landscape (Macadangdang et al., 2022). Though the host may control their amount, significant accumulations of retrotransposons have been tolerated during evolution (Lander et al., 2001; Nishihara et al., 2006). Currently, 42% of the mammalian genome is derived from retrotransposons, including ancient long terminal repeat (LTR) and non-LTR retrotransposons (Cordaux and Batzer, 2009; Fujiwara, 2015; Lander et al., 2001). The non-LTR retrotransposons, like the short-interspersed elements (SINEs) and long-terminal interspersed elements (LINEs), are highly abundant and occupy about 30% of mammalian genomes, but the majority of them are kept silent to maintain genome stability (Cordaux and Batzer, 2009). Examples of active and well-known non-LTR retrotransposons are group II introns in bacteria and R2 elements in arthropods (Eickbush and Eickbush, 2015; Stamos et al., 2017; Toor et al., 2010). The bacterial Group II introns often contain an intron RNA with a well-folded tertiary structure and intron-encoded protein (IEP) with endonuclease and reverse transcriptase activities (Chung et al., 2022; Haack et al., 2019). The intron RNA forms an RNP complex with IEP protein to recognize the DNA target via the exon binding site (EBS) sequence (Guo et al., 1997). Further, the DNA sense strand was cleaved by the RNP complex and linked to the intron RNA covalently. Following the cleavage of antisense strand DNA, the IEP protein synthesized the cDNA of intron RNA for duplication at a new site (Chan et al., 2018; Manigrasso et al., 2020; Marcia and Pyle, 2012). Based on understanding its molecular mechanism of DNA targeting and duplication, Group II introns have been repurposed as sequence-specific gene manipulation tools, such as Targetron (Mohr et al., 2013; Velázquez et al., 2022).

The eukaryotic R2 also uses the target primed reverse transcription (TPRT) process for retrotransposition via the R2 protein and less-structured RNA template and only resides in 28S DNA due to its high specificity of DNA-locus recognition (Burke et al., 1987; Luan and Eickbush, 1995). Two regulatory regions within the R2 RNA, 3’-UTR (249 nt, hereafter 3’-RNA) and 5’-ORF (323 nt, hereafter 5’-RNA), have been identified to associate with R2 protein and regulate the TPRT process (Eickbush and Eickbush, 2015; Jamburuthugoda and Eickbush, 2014). However, the molecular details about how R2 machinery recognizes and cleaves the ribosomal DNA and the regulatory roles of R2 RNA are largely unknown. A comprehensive exploration of the R2 machinery will help to understand the molecular evolution of highly active retrotransposons from prokaryotes to lower eukaryotes and shed light on the retrotransposition mechanism of poorly understood and silent retrotransposons in higher eukaryotes, like human LINE-1 which occupies 17% of the human genome (Lander et al., 2001). DNA-targeting details are also critical for developing retrotransposon-based molecular tools, especially gene-insertion tools.

Here we determined the cryo-EM structures of *Bombyx mori* retrotransposon complexes of R2 protein with 3’-RNA, 5’-RNA, or full-length mimic RNA (L-RNA), with or without ribosomal DNA at an overall resolution of 3.0 Å to 3.7 Å. Validated by sequencing analysis, we revealed two essential DNA regions that can be recognized by R2 retrotransposon. The R2 protein recognizes the DNA-recognition region (Drr) and unwinds the DNA-cleavage region (Dcr) which contains the cleavage site. Biochemical analysis suggests that the association of 3’-RNA accelerates the first-strand DNA cleavage by R2 protein and initiates the reverse transcription starting from the 3’ tail, as observed in the structure. In the 3’-RNA bound complex, the second-strand DNA cleavage is strongly inhibited, which supervises the sequential and safe process of TPRT. Removing 3’-RNA by reverse transcription allows 5’-RNA to functionally interact with the R2 protein and activates the second-strand DNA cleavage. Our results revealed the stepwise retrotransposition mechanism within the R2 machinery and the DNA targeting details that may provide practical insights for gene-editing reprogramming.

## Results

### DNA targeting specificity by R2 protein

Since DNA recognition is the first step for target insertion by retrotransposon (Figure 1A) (Jamburuthugoda and Eickbush, 2014), we first explored the target specificity by R2. The reported results indicate that the R2 protein recognizes a minimum 60 bp rDNA sequence with high fidelity (Christensen and Eickbush, 2004). However, the R2 protein is relatively small (∼120 kDa) to recognize such a long DNA. Therefore, to further determine the sequence specificity, we screened the first-strand DNA cleavage activity by the R2 protein using the reported 60 bp rDNA substrates containing the 6 bp mutation windows (Figures 1B and S1A). The results showed that only two mutation regions, C7-T24 (hereafter DNA-recognition region, Drr) and C37-C48 (containing the cleavage site; hereafter DNA-cleavage region, Dcr), significantly decreased the DNA cleavage activity by R2 protein (Figures 1B and 1C). Modified DNA by preserving the Dcr and Drr but replacing the sequence of other regions can restore the cleavage activity similar to WT rDNA target (Figure S1B). Although mutation of the spacer region between Drr and Dcr did not affect cleavage activity, increasing or shortening the spacer length significantly decreased the DNA cleavage activity by R2 protein (Figure 1D). Mutation or deletion of the sequence beyond Drr, Dcr, and spacer regions (hereafter Redundant region, Rr) had no significant effect on the DNA cleavage activity (Figure S1C). These results suggested that the R2 protein may form a specific structure to target a 40 bp DNA.

**Figure 1.**
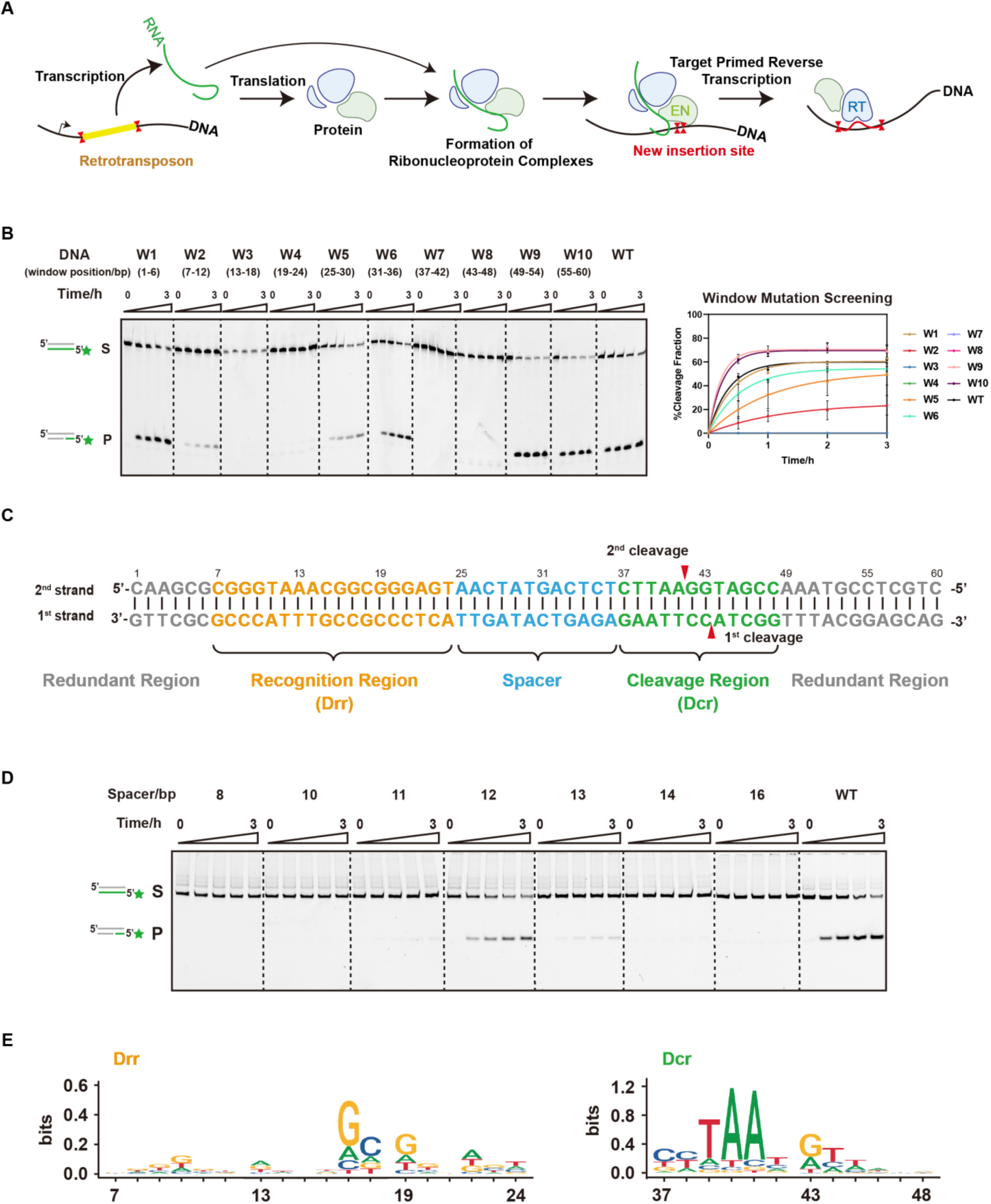
Biochemical and sequencing analysis of targeting specificity. (A) Model for the translocation mechanism of retrotransposon. The retrotransposon is transcribed into RNA and further translated into protein from DNA locus. The RNA and protein components form a complex. Then the complex reaches to a new insertion site and cleaves the DNA by protein endonuclease domain (EN). Through target primed reverse transcription by reverse transcriptase domain (RT), it inserts into the new DNA locus. (B) First-strand cleavage assay by R2 protein for window mutation screening. The DNA substrates are named as W1 to W10 according to the mutation positions from window 1 to 10. Left, the denaturing PAGE gel. Right, the plot of cleavage efficiency (n=3 each; means ± SD). (C) The native sequence of 60 bp ribosomal DNA. The number upper the sequence indicates the nucleotide position. The characters are colored according to the different regions. Gray, redundant region. Orange, recognition region (Drr). Blue, spacer. Green, cleavage region (Dcr). Red triangle indicates the cleavage sites for the first strand and second strand. (D) First-strand cleavage assay by R2 protein for spacer length variation. The number labeled for each group indicates the spacer length. (E) The deep sequencing analysis for cleavage screening. Left, the nucleotide preference in Drr. Right, the nucleotide preference in Dcr. The number blow the axis indicates the nucleotide position according to panel C (WT, wild-type DNA. S, substrates. P, products. Green colored pentagram indicates Cy5 labeling at 5’ end of DNA).

To extensively reveal the sequence preference and essential bases within Drr and Dcr, we performed the cleavage screening using the target DNA library containing 6 bp randomized windows (Figure S1D). Deep sequencing analysis indicated that R2 protein prefers to cleave DNA targets with GC-rich Drr and AT-rich Dcr (Figure 1E). Reprogramming the dsDNA by retaining these essential bases but mutating all other ones completely restored the cleavage activity as on the WT rDNA by R2 protein (Figure S1E).

### Regulatory roles of 3’-RNA and 5’-RNA on DNA cleavage

The 3’-RNA and 5’-RNA within R2 RNA are reported to associate with R2 protein and affect its activity, but the exact effects remain unclear (Christensen et al., 2006; Luan and Eickbush, 1995). Our biochemical results indicated that R2 protein alone can cleave the first strand but not the second strand of dsDNA target, and the first-strand cleavage is significantly accelerated by 3’-RNA (Figures 2A and 2B). 5’-RNA is essential to activate the second-strand cleavage by R2, while 3’-RNA can completely block this process (Figure 2B). A moderate inhibition on first-strand cleavage by 5’-RNA is also observed (Figure 2A). Of note, additions of both 3’- and 5’-RNA or 3’-RNA only into the cleavage reactions showed similar effects on R2 protein, which accelerated first-strand cleavage, and entirely abolished second-strand cleavage (Figures 2A and 2B). This result demonstrated a more dominated regulation role by 3’-RNA compared to 5’-RNA. In conclusion, R2 protein can independently recognize a 40 bp dsDNA substrate with multiple essential bases and initiate the DNA cleavage. 3’-RNA differentially promotes the first-strand cleavage and inhibit the second-strand cleavage, while 5’-RNA specifically activates the second-strand cleavage (Figure 2C).

**Figure 2.**
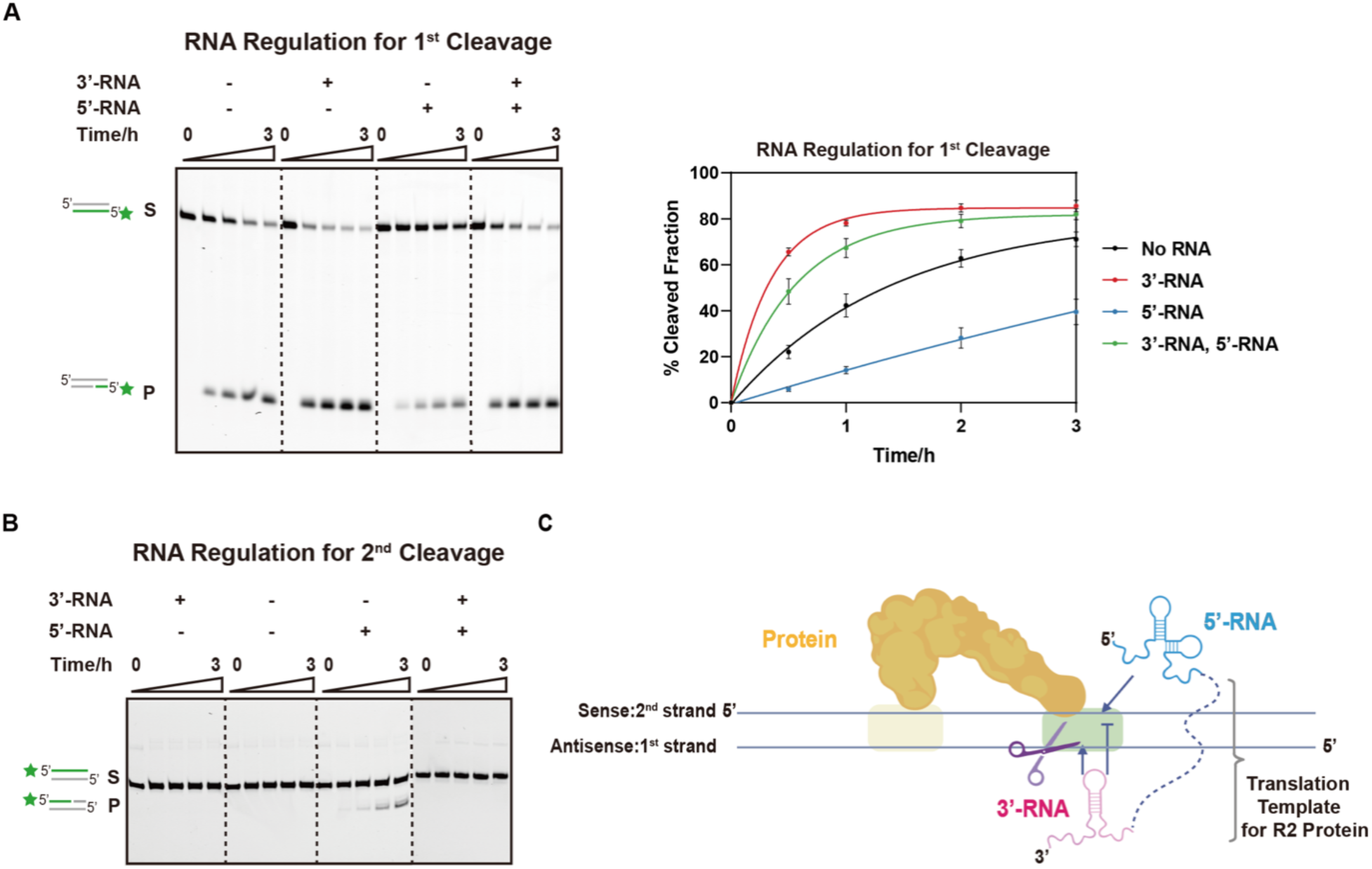
Biochemical analysis of RNA regulation on DNA cleavage. (A) First-strand cleavage assay for RNA regulation. Left, the denaturing PAGE gel. Right, the plot of cleavage efficiency (n=3 each; means ± SD). (B) Second-strand cleavage assay for RNA regulation. (C) Model for RNA regulatory roles on DNA cleavage. The protein is colored wheat. The 5’-RNA is colored blue. The 3’-RNA is colored pink. The scissor indicates the DNA cleavage. The yellow box indicates Drr on DNA. The green box indicates Dcr on DNA (WT, wild-type DNA. S, substrates. P, products. Green colored pentagram indicates Cy5 labeling at 5’ end of DNA).

### Molecular details for DNA recognition

To explore the molecular mechanism of specific DNA recognition by R2 machinery, we first reconstituted the binary complex with deactivated R2 protein (D628Y, D996A) (Christensen and Eickbush, 2005; Yang et al., 1999) and the 60 bp rDNA target (Figure S2A). Although the biochemical results indicated that they form a stable complex, cryo-EM analysis only revealed the low-resolution (∼20 Å) 2D and 3D maps, indicating the internal structural flexibility within the binary complex (Figures S2A-S2C). Given the dominant effect on the DNA cleavage by 3’-RNA, we speculated that 3’-RNA might enhance the stability of DNA and R2 protein complex. As expected, we successfully reconstituted a stable 3’-RNA bound complex containing deactivated R2 protein, rDNA, and 3’-RNA (Figure S2D) and obtained the cryo-EM map at 3.6 Å resolution for model building (Figures 3A-3C and S2E-S2I; Video S1). In general, the R2 protein consists of five domains from the N-terminus to the C-terminus: DNA-recognition region binding domain (DrrB, containing a Zinc finger and a Myb motif), RNA-binding domain (RB), reverse transcriptase domain (RT), and DNA-cleavage region binding domain (DcrB) and endonuclease domain (EN) (Figures 3A and 3C). Overall, the R2 protein displays a question mark architecture, with the RT domain showing a classic right-hand shape (Figures 3D and S3A) (Stamos et al., 2017).

**Figure 3.**
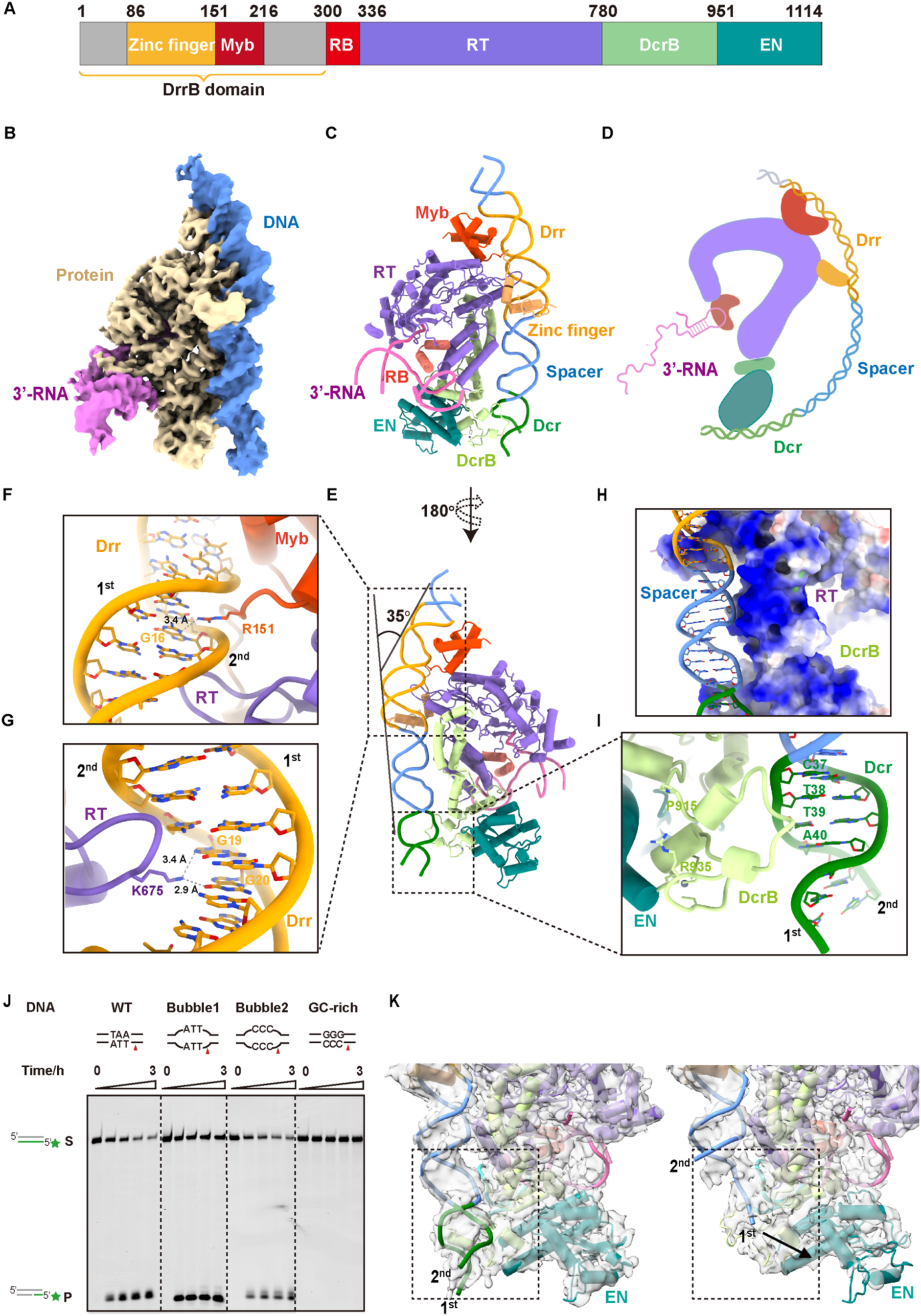
Structural analysis of DNA recognition and unwinding. (A) The domain structure of R2 protein. The number indicates the position of amino acids. DrrB domain, DNA-recognition region binding domain. RB, RNA binding domain. RT, reverse transcriptase domain. DcrB domain, DNA-cleavage region binding domain. EN, endonuclease domain. (B) The cryo-EM map of 3’-RNA bound complex reconstituted by deactivated R2 protein, DNA and 3’-RNA. The density of protein, DNA and 3’-RNA are filled in different color. (C) The atomic model for 3’-RNA bound complex. The protein domains are colored referring to panel A. The Drr on DNA is colored yellow. The spacer region on DNA is colored blue. The Dcr on DNA is colored green. 3’-RNA is colored pink. The front and back views are shown in panel C and E. (D) A cartoon model for the question mark architecture of R2 protein as well as the schematic diagram of DNA and RNA binding. The protein domains, DNA and 3’-RNA are colored accordingly. (E) The back view of the atomic model for 3’-RNA bound complex. The DNA bending angle and degree are shown. (**F** and **G**) The structure details for the interaction between DrrB domain and Drr on DNA. The amino acids involving for dG(16) (**F**), dG(19) and dG(20) (**G**) recognition are labeled. The key hydrogen bonds are shown by dashed lines and labeled with atom distance. (H) The surface charges of DcrB domain. Blue indicates a positive charge while red indicates a negative charge. (I) The structure details for the interaction between DcrB domain and Dcr on DNA. (J) First-strand cleavage assay by R2 protein for bubble DNA. S, substrates. P, products. Green colored pentagram indicates Cy5 labeling at 5’ end of DNA. Red triangle indicated the cleavage sites for the first strand. (K) Two conformations classified from 3D-classification of R2-DNA-3’-RNA ternary complex. Left, the state of pre-cleavage. Right, the state of first-strand cleavage. The DNA is unwound and the first strand towards EN domain. The arrow indicates the extension.

Referring to the atomic model, the rDNA primarily interacts with DrrB and DcrB domains (Figures 3C and 3E). Specifically, DrrB uses Myb motif and Zinc finger to capture the Drr of rDNA via electrostatic interaction and base-specific recognition (Figures 3E-3G, S3B, and S3C). The Drr also exhibits approximately 35-degree bending due to DrrB’s binding (Figure 3E). The hydrogen-bond interactions for R151 (in Myb motif) with G16 as well as K675 (in RT domain) with G19 and G20 on the second DNA strand were observed, which is consistent with the sequencing data showing the G base preferences at these positions (Figures 1E, 3F, and 3G). Following, the DNA spacer region interacts with the DcrB through weak electrostatic interactions, and it still maintains the B-form conformation (Figure 3H). Within the Dcr, we also observed the interaction interface between the DNA C37-A40 (on the second strand) and the P915-R935 motif in the DcrB domain (Figure 3I). Of note, Dcr remains in the double-stranded form and is far from the EN active core, suggesting a structural state of DNA recognition before cleavage (Figures 3C and 3E).

As specific recognition between DcrB and the A or T base in Dcr was not observed, we speculated that the preference for AT bases is due to the fact that AT-rich dsDNA is more easily to be unwound for cleavage. Indeed, dsDNAs containing GC-rich or mutated AT bubble in Dcr can be efficiently cleaved by R2 protein compared to WT rDNA (Figure 3J). 3D classification also revealed a conformation in which the Dcr is unwound and the first DNA strand towards the EN domain, corresponding to the first-strand cleavage state (Figure 3K). Conformation with second-strand pointing to the EN domain was not observed even by extensive 3D classification, indicating that while 3’-RNA greatly enhances the stability of DNA-R2 protein interaction and fixes the DNA-binding orientation on the R2 protein, it also limits the structural flexibility required for loading the second strand to EN catalytic core. Overall, R2 protein recognizes the GC-rich Drr and unwinds the AT-rich Dcr to initiate the dsDNA cleavage process.

### Essential segments within 3’-RNA regulate the R2 activity

Combining icSHAPE analysis and reported secondary structure of 3’-RNA(Moss et al., 2011; Ruschak et al., 2004), we located two 3’-RNA stems (corresponding to S1 and S5 in the secondary structure) in the cryo-EM map (Figures 3B, 4A, 4B and S4A; Video S1). The structure also revealed hydrogen-bond interaction between the U131 and R311 (in RB domain) (Figure 4C). Moreover, the 3’-RNA polyA tail was clearly observed and pointed to the RT catalytic pocket (Figure 4D), indicating a TPRT initiation state in which cDNA will be reverse transcribed starting from the 3’-RNA polyA tail. Of note, shortening the length of polyA tail significantly decreased the TPRT activity (Figure S4B). Likely, other regions of the 3’-RNA don’t have stable 3D structure, so we are not able to observe them in the cryo-EM map, as well as the direct interaction between 3’-RNA and DNA.

**Figure 4.**
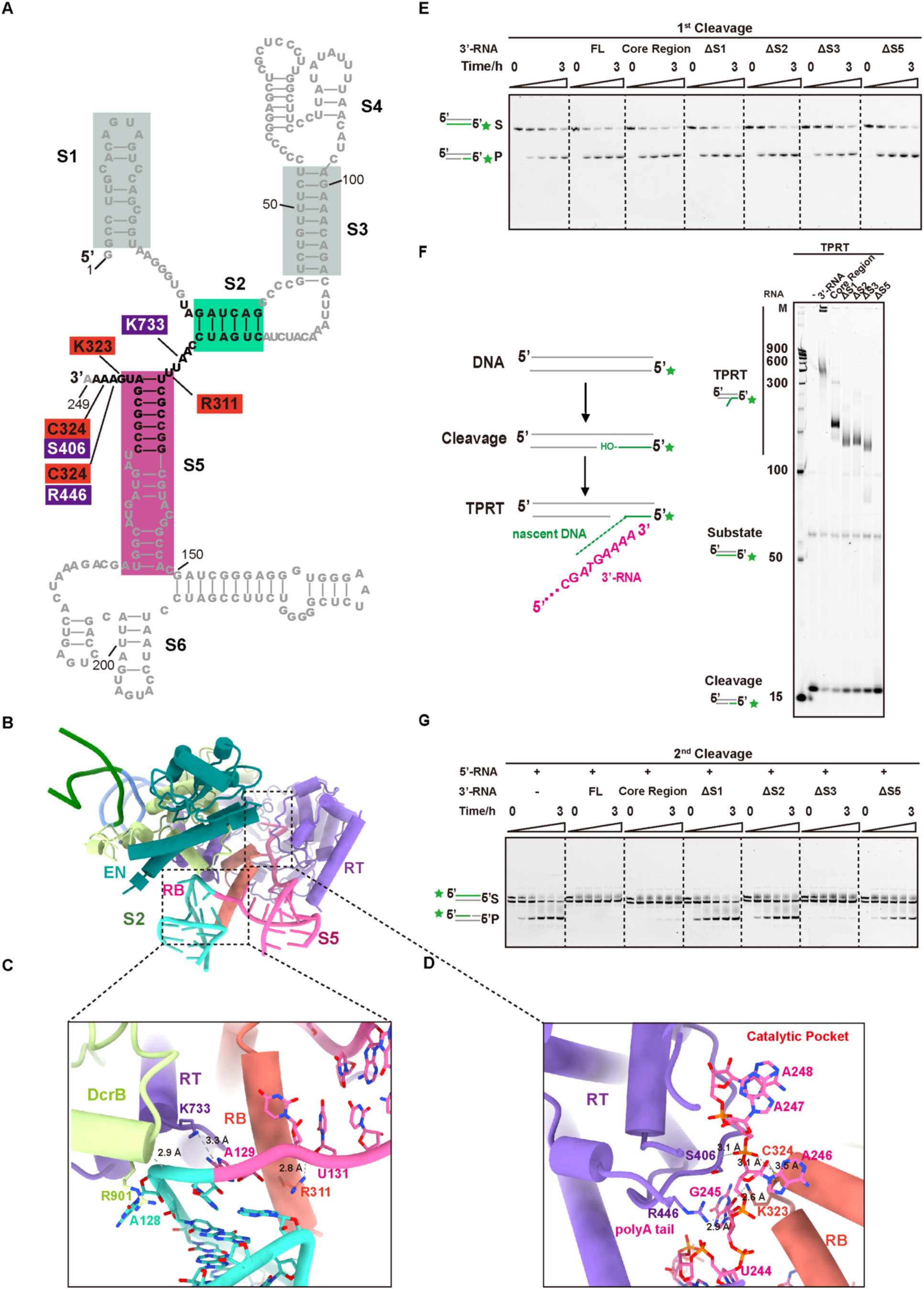
Structural and biochemical analysis of 3’-RNA regulation. (A) The secondary structure of 3’-RNA. The ribonucleotides which were identified in cryo-EM map are shown in black, while others are shown in gray. The shadows indicate the segment S1, S2, S3, and S5. The S2 is colored turquoise. The S1 and S3 are colored gray. The S5 is colored pink. The interaction pairs between ribonucleotides and residues in RB domain (red) and RT domain (purple) are linked with solid lines. (B) S2 is colored turquoise and S5 is colored pink. The protein domains are colored according to Figure 3A (C) The structure details for the interaction between R2 protein and 3’-RNA. The hydrogen-bond interactions are shown by dashed lines and labeled with atom distance. (D) The structure details of the 3’-RNA tail pointing to catalytic pocket of RT domain. The hydrogen-bond interactions are shown by dashed lines and labeled with atom distance. (E) First-strand cleavage for truncated 3’-RNAs. The core region means that removing S4 and S6 from the full length 3’-RNA. ΔS1, ΔS2, ΔS3, and ΔS5 mean that removing S1, S2, S3, and S5 from the core region, respectively. (F) Target primed reverse transcription (TPRT) assay for truncated 3’-RNA. The reaction time is 16 hours. Left, the schematic diagram of TPRT. After cleavage, the RT domain of R2 protein catalyzes the reverse transcription using 3’-OH as primer and RNA as template. Right, the denaturing PAGE gel. (G) Second-strand cleavage for truncated 3’-RNA. (S, substrates. P, products. Green colored pentagram indicates Cy5 labeling at 5’ end of DNA.)

To further dissect the regulation effects on R2 activity (DNA cleavage and TPRT) by the different regions of 3’-RNA, especially the structurally invisible regions, we divided the 3’-RNA into six segments (S1 to S6) referring to our refined secondary structure model (Figure 4A). Of note, the removal of S4 and S6 had minimal impact to the regulation effects on R2 activity by 3’-RNA, including accelerating first-strand cleavage, initiating TPRT, and blocking the second-strand cleavage (Figures 4E-4G, S4C, and S4D). Therefore, we identified an essential core within the 3’-RNA including S1, S2, S3 and S5. Within the 3’-RNA core, removing S3 eliminates its promoting effect on the first-strand cleavage and decreased its inhibition on the second-strand cleavage, removing S5 eliminates its ability to activate TPRT, and removing both S1 and S2 eliminates its inhibition on the second-strand cleavage (Figures 4E-4G, S4C, and S4D). In conclusion, by combining structural and biochemical analysis, we have uncovered that the 3’-RNA exhibits multi-regulatory effects on R2 activity using different functional segments.

### 5’-RNA displays a “three-claw” structure to capture R2 protein

To explore the activation mechanism of second-strand cleavage by 5’-RNA, we reconstituted 5’-RNA bound complex containing deactivated R2 protein, 5’-RNA, and the 60 bp rDNA for structural analysis, and obtained a cryo-EM map at 3.5 Å resolution (Figures 5A and S5; Video S1). The majority of the 5’-RNA density was visible, which displayed a “three-claw” platform to securely capture R2 protein (Figure 5A). Combining icSHAPE analysis and reported secondary structure (Hart et al., 2008; Kierzek et al., 2008; Moss et al., 2011), we built the nearly complete atomic model of 5’-RNA (Figures 5B, 5C, and S6). Claw1 is the largest and most complicated one (U23-C124), which consists of element I and element II (pseudoknot region) and forms a large electrostatic interaction interface with the RT domain and DcrB domain (Figures 5B, 5C, and S6A). Claw2 is relatively simple and only consists of element III (G125-U165), which binds to the back of the RT catalytic pocket (Figures 5 and S6B). Claw3 consists of elements IV to VI, which interact with RB domain and the non-catalytic region of EN domain, respectively (Figures 5B, 5C, and S6C). A further dissection of the three 5’-RNA claws indicates that claw3 is essential and sufficient for second-strand cleavage activation, and claws 1 and 2 play the auxiliary effects (Figure S7).

**Figure 5.**
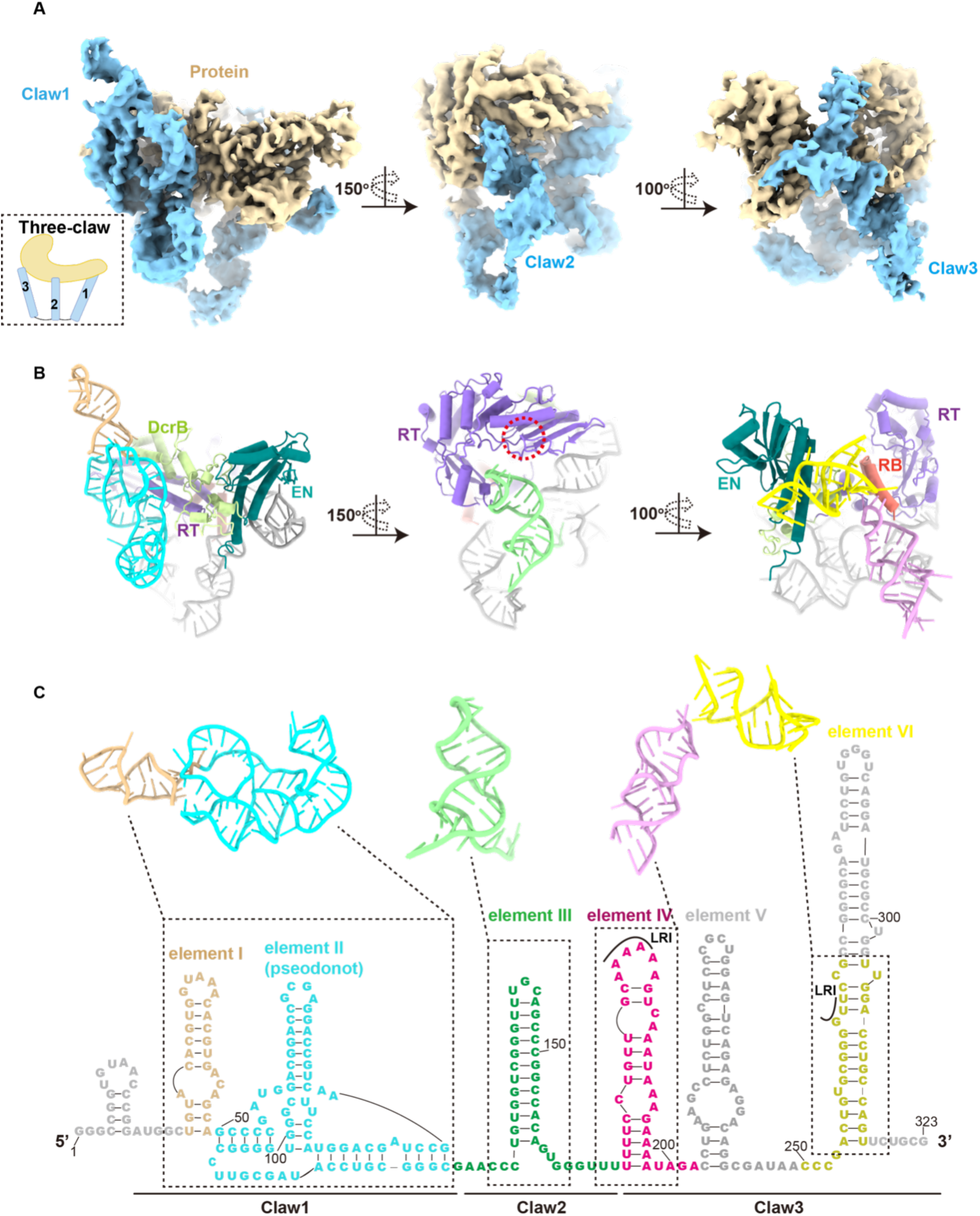
Structural analysis of 5’-RNA bound complex. (A) The cryo-EM map of 5’-RNA bound complex reconstituted by deactivated R2 protein, rDNA and 5’-RNA. The cartoon model shows the architecture of protein hold by “three-claw” 5’-RNA. The density of protein is colored in wheat. Claws of 5’-RNA is colored blue. (B) The atomic model for 5’-RNA bound complex. The protein domains are colored according to Figure 3C. The element I is colored wood, element II is colored cyan, element III is colored green, element IV is colored plum, and element VI is colored yellow. The red dashed circle means the catalytic pocket. (C) The secondary structure of 5’-RNA. Ribonucleotides of 5’-RNA that had not been observed are colored gray. Element I-VI are colored according to panel B. The long-range interaction (LRI) between element IV and element VI is labeled with solid lines.

Though our biochemical results strongly suggested that rDNA, 5’-RNA, and R2 protein can form a stable ternary complex, the density of rDNA and DrrB domain was not observed in this EM map due to structural flexibility (Figures 5A and S5A). We further confirmed this via EM map comparison to the reconstituted binary complex of 5’-RNA and R2 protein (Figures S8A and S8B). Structural comparison between 3’-RNA and 5’-RNA bound complexes shows that the Drr and 5’-RNA claw1 share a similar binding interface on R2 protein (Figure S8C). Thereby, 5’-RNA claw1 may compete the DNA-protein interaction and dissociate the Dcr from R2 protein (Figure S8C). Whereas, the Drr still interacts with DrrB, forming a DrrB-DNA complex and waving around the R2 stable core (Figure S8C). With low structural complexity, DrrB can form the phase condensates *in vitro*, indicating that it may undergo multiple conformations to serve as a flexible but faithful anchor for capturing DNA substrate via Drr interaction (Figure S8D). Of note, this flexible DNA-binding state may allow the DNA to adjust its angle and deliver the second strand into the EN catalytic pocket, thereby activating the second-strand cleavage (Figure S8C). Moreover, EN domain also adopts a 12-angstrom shift due to 5’-RNA association, which may help EN domain better capture the second-strand DNA (Figure S8E; Video S1). However, this hypothesis that the flexible DNA-binding state allows second-strand cleavage needs to be further proved in future work. Overall, the 5’-RNA forms a complicated three-claw structure to capture the non-catalytic region of R2 protein and compete with the interaction interface between R2 protein and Dcr within rDNA.

### The sequential retrotransposition by R2 machinery

The 5’-RNA, besides protein binding competition with the Dcr, its claw3 binding site overlaps with the 3’-RNA binding site on RB domain (Figure 6A). In order to further illustrate how 3’- and 5’-RNA coordinate to regulate R2 machinery when co-existing, we synthesized the full-length R2 RNA mimicking containing both 3’-RNA, 5’-RNA, and a 20 bp linker (hereafter L-RNA) and obtained the cryo-EM structure of the L-RNA bound complex reconstituted by deactivated R2, L-RNA, and rDNA at 3.0 Å resolution (Figure S9; Video S2). Within the L-RNA bound complex, 3’-RNA still binds to the RB domain with polyA tail pointing to RT catalytic core suggesting the TRPR initiation state as 3’-RNA bound complex, and the claw3 of 5’-RNA dissociates from the protein due to 3’-RNA competition (Figures 6A and S10A). Of note, the claw1 shows an identical binding pattern with R2 protein compared to 5’-RNA bound complex, and about 15-degree displacement of claw2 occurs due to the claw3 dissociation (Figure S10B). This analysis suggests, even under 5’-RNA competition, 3’-RNA can still stably interact with R2 protein, thereby dominating the regulation effect. In consistent, our biochemical results indicate that the second-strand cleavage is significantly inhibited when 3’-RNA and 5’-RNA exist simultaneously (Figures 2B and 6B). However, if the cleavage reaction is supplemented with dNTPs to initiate the TPRT which unfolds the 3’-RNA structure, the inhibition effect on the second-strand cleavage by 3’-RNA can be relieved (Figure 6B). In R2 machinery, the TPRT uses DNA primer dispensing with a complementary sequence to the RNA template (Bibillo and Eickbush, 2004; Jamburuthugoda and Eickbush, 2014). Thereby, blocking the second (antisense) strand cleavage by 3’-RNA can ensure that only primer produced by the first (sense) strand cleavage is available, but not the second strand, to synthesize the R2 cDNA, thus ensuring correctly orientated insertion into the new site. Of note, inverted insertion is fatal for R2, which relies on the 28S promoter for expression (Eickbush and Eickbush, 2010; Eickbush et al., 2008).

**Figure 6.**
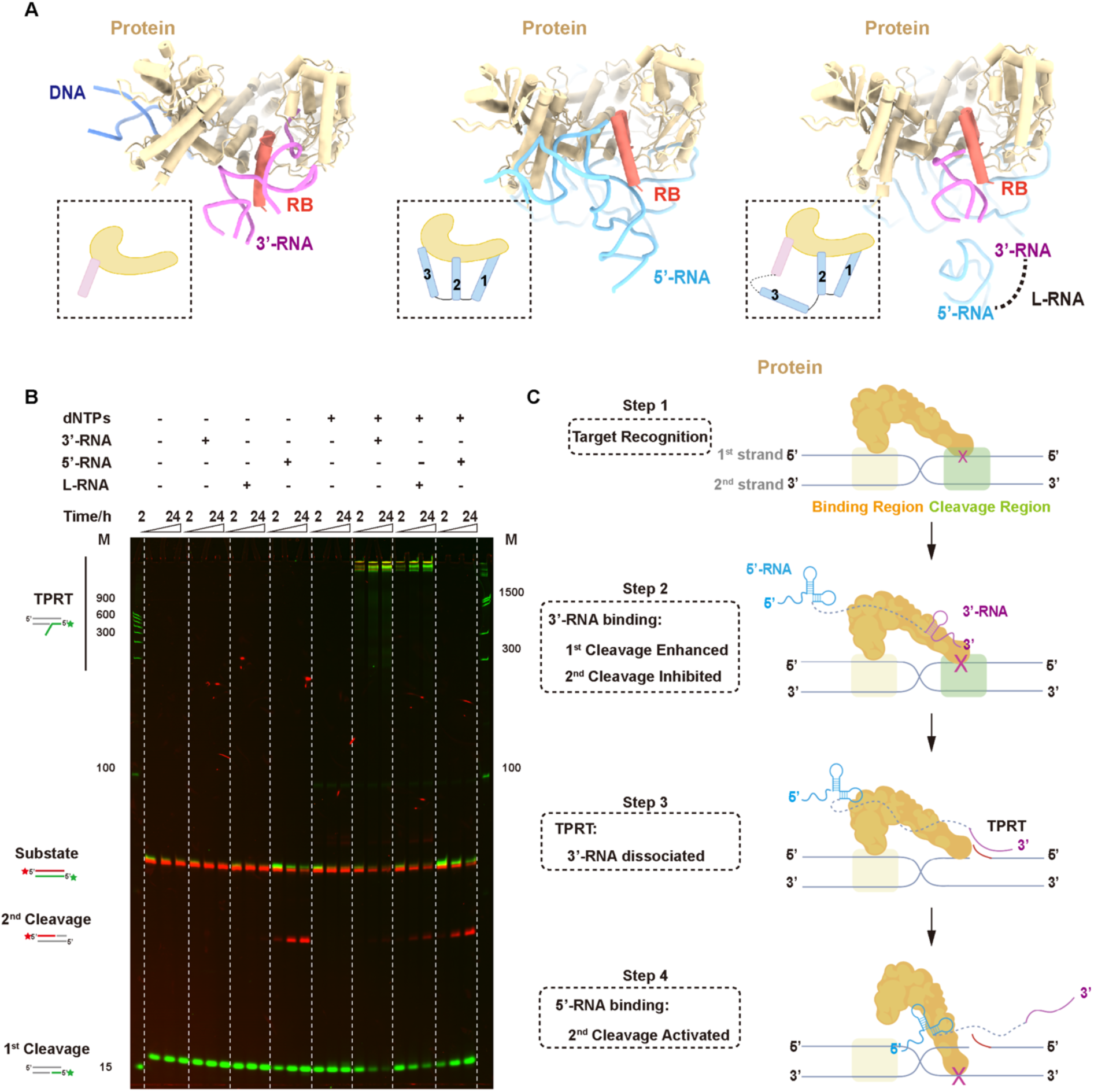
Stepwise retrotransposition regulated by R2 RNA. (A) Atomic model comparison of 3’-RNA (left), 5’-RNA (middle), and L-RNA (right) bound complexes. R2 protein is colored wheat, and RB domain is colored red. DNA, 3’-RNA and 5’-RNA are colored dark blue, pink and light blue, respectively. Cartoons illustrating protein and RNA interactions are shown. (B) Cleavage and target primed reverse transcription (TPRT) assay for RNAs. The reaction time is 24 hours. Left, the schematic diagram of TPRT. The first strand is labeled by Cy5 (green), the second strand is labeled by FAM (red). Fluorescence channels are merged via ImageJ. (C) Stepwise working model of R2 retrotransposon. The protein is colored wheat. The 5’-RNA is colored blue. The 3’-RNA is colored pink. The cross indicates the DNA cleavage. The yellow box indicates Drr on DNA. The green box indicates Dcr on DNA.

In conclusion, we propose a stepwise model for the retrotransposition within R2 machinery: Step 1, R2 protein recognizes the Drr sequence on the rDNA substrate and initiates the first-strand cleavage in the Dcr; Step 2, R2 3’-RNA preferentially associates with RB domain and RT catalytic core and tightens the DNA-protein interaction, accelerating the first-strand cleavage, suppressing the second-strand cleavage, and serving as the template to initiate TRPT from the 3’ tail; Step 3, as TPRT proceeds, the structure of 3’-RNA is unfolded, causing it to dissociate from R2 protein; Step 4, the dissociation of 3’-RNA enables 5’-RNA to functionally interact with R2 via a three-claw way and loosen the DNA to adjust the conformation for second-strand DNA cleavage (Figure 6C).

## Discussion

### The decreased structural and functional complexity of RNA components in more evolved retrotransposons

Using cryo-EM and biochemistry analysis, we revealed the mechanism of target recognition and sequential regulation by structural RNAs in eukaryotic R2 retrotransposition. The results showed that the two regulatory RNAs form complicated tertiary structures and bind to R2 proteins, respectively, to regulate the DNA cleavage and TPRT activity, ensuring the safe and sequential progress of retrotransposition. The flexible N-terminal domain DrrB in the R2 protein folds into two DNA-binding motifs on the protein surface to recognize and anchor the DNA substrate. This is utterly different from Group II intron machinery, the hypothetical ancestor of R2 in prokaryotes, which uses the RNA EBS region to recognize complementary DNA (Eickbush, 1999; Guo et al., 1997). Moreover, compared to Group II intron RNA, the R2 RNAs have much simpler structures and lost the self-splicing activity. These differences suggest that, from prokaryotes to eukaryotes, the structural complexity of RNA components in the retrotransposon machinery is gradually diluted, and its catalytic activity and function of DNA recognition are largely replaced by protein enzymes (Figure S10C). Consistently, in the retrotransposon of even higher organisms, like human LINE-1, the RNA component has been completely transformed into a linear template similar to mRNA, without complicated structure or catalytic activity (Figure S10C) (Grechishnikova and Poptsova, 2016). Thereby, what positive effects this evolutionary trend may play on lives deserves more study. It is also an interesting question whether non-structured, linearized RNA components allow the host to better control the activity of retrotransposon and thus ensure the stability of the genome, as seen for human LINE-1.

### Limited knowledge about RNA structure and function

Although 5’-RNA forms more complex structures and interaction interfaces with R2 protein than 3’-RNA, our biochemical results suggested that 3’-RNA shows a more prominent effect on R2 activity. Structural studies have also demonstrated that the association of 3’-RNA and the critical RNA-binding region in the R2 protein cannot be competed by 5’-RNA. Only by reverse transcription within TPRT process, the 3’-RNA can be structurally unfolded and released, allowing the complete and functional association of 5’-RNA with the R2 protein. On the one hand, this analysis suggests that our current understanding of the connection between RNA structure and function is still minimal. On the other hand, it implies that we may design smaller but highly efficient functional RNA molecules for regulating the function of proteins, similar to the RNA aptamers often obtained via screening (Lee et al., 2023).

### R2 machinery provides an engineerable gene-editing platform

Recently, R2 has been used for gene insertion in eukaryotic cells with the conserved insertion site and sequence (Kuroki-Kami et al., 2019; Su et al., 2019). Our results offer timely information for engineering and improving the R2-based gene-editing tools. For example, to reprogram the insertion template, we can preserve the essential 3’-RNA and 5’-RNA elements, and use the designed sequence to replace the junction RNA region as the reverse transcription template. Our results also suggest that *Bombyx mori* R2 shows a certain level of sequence tolerance (Figures 1B and 1E), which offers some flexibility for target site choices. To precisely reprogram the target sites in future studies, comprehensive engineering or replacement of the N-terminal DrrB domain of R2 protein should be performed. Noteworthy, the R2 family has an abundant diversity of homologous species using different N-terminus to recognize their preferred DNA sequences (Kojima and Fujiwara, 2005; Luchetti and Mantovani, 2013), which may provide various options for the target sites.

## Acknowledgements

EM data were collected at the Tsinghua Cryo-EM facility. The data were analyzed using the Bio-Computation platform at the Tsinghua University Branch of the Chinese National Center for Protein Sciences (Beijing). We thank J.L. Lei, X.M. Li and X.D. Li for expert electron microscopy assistance. We thank T. Yang, Y.K. Wang, A.B. Jia for computational support. We thank M.E. Wilkinson and F. Zhang for their kind advice on the model building.

## Author contributions

J.J.G.L supervised the project. J.J.G.L, J.W, P.D and S.Q.T designed the experiments. P.D, S.Q.T, Q.Y.Y purified the proteins and performed the biochemical assays and analysis. Z.B.B and Y.L performed phase separation assays and analysis. L.S and Q.C.Z. did the icSHAPE analysis. J.W, P.D, S.Q.T and H.Z.Z did the structural analysis and built the atomic model. J.J.G.L, J.W, P.D and S.Q.T wrote the manuscript with the help from all authors.

## STAR★METHODS

### Key resources table

All the reagents used in this work are commercially available. All the protocols have been described in detail in the text. All the software used in this work have been indicated with references and are available for academic usage.

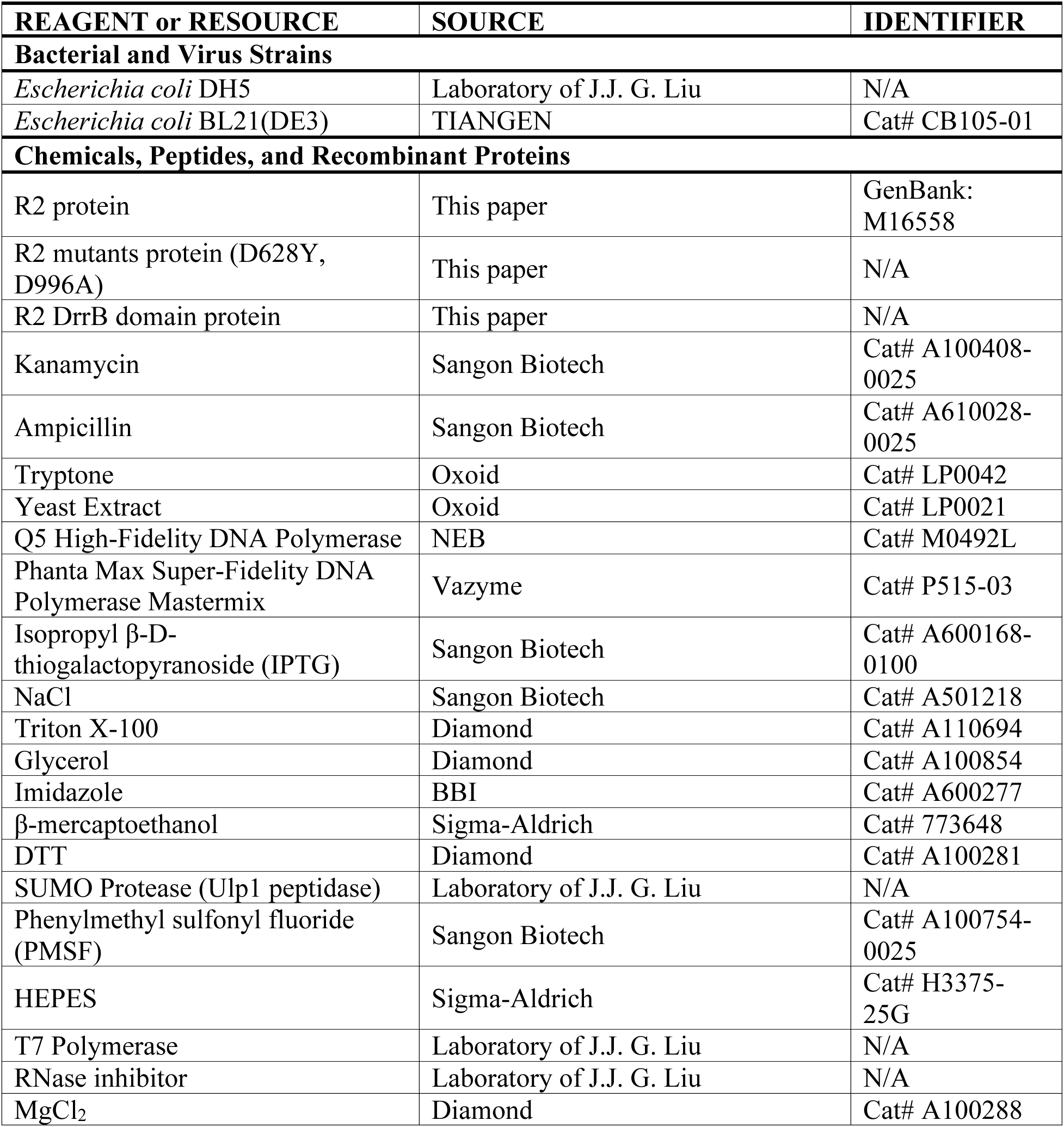
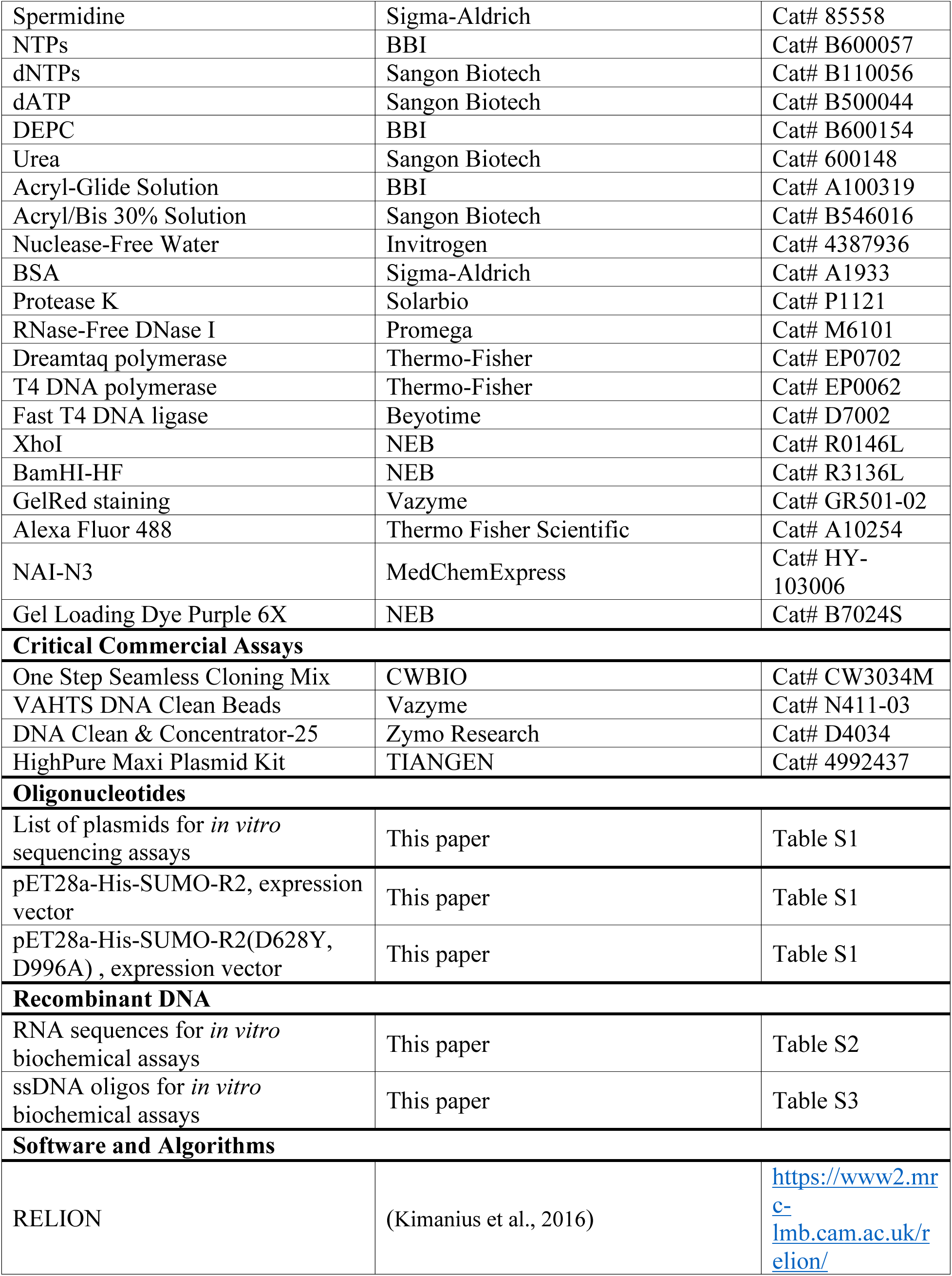
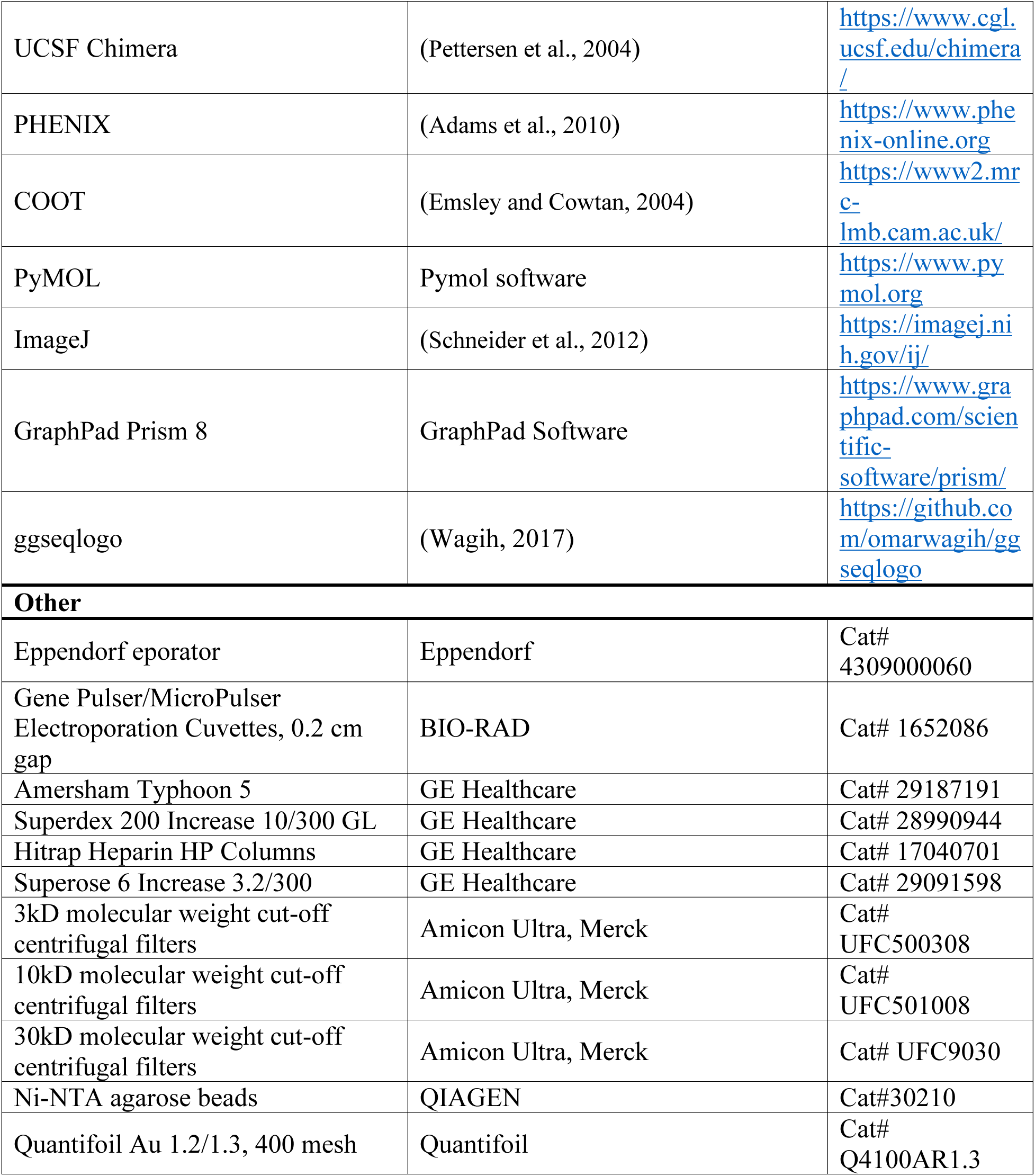

### Resource availability

#### Lead contact

Readers are welcome to comment on the online version of the paper. Correspondence and requests for materials should be addressed to the lead contact J.J.G.L.

#### Materials availability

Plasmids generated in this study are listed in Table S1 and will be deposited to Addgene or are available upon request.

#### Data and code availability

The electron density maps have been deposited to the Electron Microscopy Data Bank (EMDB) under the accession numbers of EMD-35347, EMD-35348, EMD-35349, and EMD-35350 which are publicly available as of the data of publication. The atomic coordinates and structure factors have been deposited to the Protein Data Bank (PDB) under the accession number of 8IBW, 8IBX, 8IBY, and 8IBZ which are publicly available as of the data of publication. The raw cryo-EM micrographs and movies used in this study will be shared by corresponding author upon request. Any additional information required to re-analyze the data reported in this paper is available from the corresponding author upon request.

### Method details

#### Plasmids construction

The sequence of *Bombyx mori* R2 gene was downloaded from NCBI. Bacteria- and insect-codon optimized *Bombyx mori* R2 gene was ordered from Sangon Biotech. R2 gene was cloned into pET28a-based vector with an N-terminal 10xHis-SUMO tag by homologous recombination (One Step Seamless Cloning Mix, CWBIO). For D628Y and D996A mutations in RT domain and EN domain of R2, mutagenetic PCR was performed to amplify mutated fragment and cloned into pET28a-based vector by homologous recombination. DrrB domain of R2 protein was amplified from R2 gene and cloned to pET28a-based vector. To construct the cleavage screening library, ribosomal DNA containing 6 bp randomized sequence in different windows were ordered from RuiBiotech and cloned into pUC57 backbone via homologous recombination. The plasmids and related description are listed in Table S1.

#### Protein expression and purification

The wide-type R2 protein was purified for *in vitro* biochemical assay and the deactivated R2 protein was purified for structural analysis. The plasmid was transformed into BL21 (DE3) *Escherichia coli* cells (TIANGEN) and incubated overnight at 37 °C first in 100 mL start-culture LB medium with 50 μg/ml Kanamycin. It was transferred to a large 1L culture with 50 μg/ml Kanamycin and grown at 37 °C to an A_600_ of 1.2. Protein expression was induced at 20 °C for 20 hours with 0.2 mM IPTG. Cells were harvested and resuspended in lysis buffer (20 mM HEPES-Na pH7.5, 600 mM NaCl, 40 mM Imidazole, 10% Glycerol, 1% Triton, 1 mM PMSF and 5 mM β-mercaptoethanol). Cells were lysed by sonication and the lysate was centrifuged at 15000 g for 60 minutes at 4 °C. The supernatant was loaded onto Ni-NTA gravity column and incubated. The resin was washed with wash buffer (20 mM HEPES-Na pH7.5, 600 mM NaCl, 40 mM Imidazole, 10% Glycerol, 0.1% Triton) and then eluted with elution buffer (20 mM HEPES-Na pH7.5, 400 mM NaCl, 300 mM Imidazole, 10% Glycerol, 0.01% Triton). ULP1 and 5 mM DTT were added into the eluted protein and incubated for 1 hours at 4 °C to remove the N-terminal 10xHis-SUMO tag. The protein was loaded onto 10 mL HiTrap Heparin HP column (GE Healthcare), equilibrated with heparin buffer (10 mM HEPES-Na pH7.5, 150 mM NaCl, 10% glycerol, 2 mM DTT, 0.01% Triton), and eluted with a linear gradient of heparin buffer B (10 mM HEPES-Na pH7.5, 2 M NaCl, 10% glycerol, 2 mM DTT). The peak fractions were pooled and concentrated to 1 mL using 30 kDa molecular weight cut-off centrifugal filters (Merck Millipore). The protein was stocked at - 80 °C after flash-frozen in liquid nitrogen. The DrrB domain protein was expressed and purified for in *vitro* phase separation assays. The expression process was the same as mentioned above. For purification, the lysis buffer (20 mM HEPES-Na pH7.5, 400 mM NaCl, 40 mM Imidazole), elution buffer (20 mM HEPES-Na pH7.5, 200 mM NaCl, 300 mM Imidazole, heparin buffer (10 mM HEPES-Na pH7.5, 150 mM NaCl, 2 mM DTT) and heparin buffer B (10 mM HEPES-Na pH7.5, 2 M NaCl, 2 mM DTT) were different. The protein was further purified by size exclusion chromatography column (Superdex 200 Increase 10/300, GE Healthcare) with S200 buffer (10 mM HEPES-Na pH7.5, 200 mM NaCl, 2 mM DTT).

#### RNA transcription and Purification

The DNA sequence of 3’-RNA and 5’-RNA were ordered from Sangon Biotech. The DNA sequence of truncated 3’-RNAs were assembled by overlap PCR from 3’-RNA. The DNA sequence of L-RNA was assembled by overlap PCR from 3’-RNA and 5’-RNA. The sequences were then amplified and added T7 RNA polymerase promoter by PCR. The amplification products were the template for *in vitro* transcription (IVT). The 500 uL reaction was performed at 37 °C for 4 hours in IVT buffer (30 mM Tris pH 8.1, 25 mM MgCl_2_, 0.01% Triton, 2 mM spermidine) with 4 mM NTPs and 0.05 mg/mL T7 RNA polymerase. The DNase reaction buffer and 25 u RNase-free DNase I (Promega) were added to digest the DNA template. After 1 hour digestion at 37 °C, the product was centrifuged at 14000 g for 30 minutes to remove precipitates. The RNA product was extracted by phenol chloroform and centrifuged using 10 kDa molecular weight cut-off centrifugal filters (Merck Millipore) to dissolve in DEPC water and stocked at -80 °C. The RNA sequences and related descriptions are listed in Table S2.

#### *In vitro* cleavage assays

For 60 bp native ribosomal DNA and mutant DNA, the labeled dsDNA substrates were prepared by PCR extension. The non-labeled 60 nt ssDNA as template was ordered from Tsingke Biotech. The 5’-NH_2_ modified 14 nt primer was ordered from Sangon Biotech. The modified primer was labeled via the reaction between -NH_2_ and Cy5-NHS. It was purified by ethanol precipitation. The PCR extension was performed to obtain the labeled dsDNA. Then the dsDNA was purified by DNA Clean & Concentrator-25 (Zymo Research) and diluted to nuclease-free water (Invitrogen). For 42 bp truncate ribosomal DNA, the labeled ssDNA and non-labeled ssDNA were ordered from Sangon Biotech and annealed to obtain the labeled dsDNA. For double labeled 60 bp native ribosomal DNA, the FAM labeled ssDNA and Cy5 labeled ssDNA were ordered from Sangon Biotech and annealed.

*In vitro* cleavage assays were performed in cleavage buffer (10 mM HEPES-Na 7.5, 400 mM NaCl, 5 mM MgCl_2_, 2 mM DTT, 10% Glycerol, 0.01% Triton and 0.1 mg/mL BSA) and reacted at room temperature. The ratio of R2 protein to dsDNA substrate was 50:1 and the final concentration of labeled dsDNA was 10 mM. The reaction was started by adding the labeled dsDNA into the reaction mixture. The samples were collected at the following time points: 0 h, 30 min, 1 h, 2 h, 3 h. For the cleavage assay of RNA regulation, the ratio of RNA to R2 protein was 1.5:1. R2 protein and RNA were assembled at RT for 30 minutes in cleavage buffer and cooled on ice for 5 minutes before adding the dsDNA substrate. For the TPRT assay, the final concentration of dNTP mix was 125 uM and was added at last to start the reaction. Protease K (Solarbio) was added to digest the protein and reacted at 55 °C for 15 minutes. The samples were collected at 2 h, 16 h, 24 h. All cleavage samples collected above were analyzed in 15% urea-PAGE and visualized using Amersham Typhoon 5 (GE Healthcare). The cleavage efficiency was quantified using Image J and the curves of cleavage efficiency were plotted using One-Phase-Decay model in Prism 8 (GraphPad). The RNA sequences and related description are listed in Table S3.

#### DNA cleavage screening assay

The deep sequencing workflow for cleavage screening assay was inspired and modified from previously described PAM depletion assay (Figure S1D) (Karvelis et al., 2015; Sun et al., 2023). The 60 bp native ribosomal DNA was divided into 10 windows, each contained 6 bp nucleotides. The ribosomal DNA was substituted with randomized sequences in these windows, respectively. Oligonucleotides contained the 6 bp randomized window and the ribosomal DNA were ordered from RuiBiotech and constructed in pUC57 backbone via homologous recombination. The recombination products were transformed into DH5α *Escherichia coli* (TIANGEN) and incubated overnight at 37 °C in 100 mL LB medium (100 μg/ml Ampicillin). All the culture was harvested to extract the pUC57_N6 plasmid library using HighPure Maxi Plasmid Kit (TIANGEN).

The pUC57_N6 library was cleaved by R2 protein and 5’-RNA complexes in cleavage buffer to linearize the plasmid. The ratio of 5’-RNA, R2 protein and plasmid library was 30:20:1. The plasmid library input was 10 μg. After 48 hours, the reaction was quenched with loading buffer (Gel Loading Dye Purple 6X, NEB) supplemented with 20 mM EDTA and 25 μg/mL heparin. The Protease K (Solarbio) was added to digest the protein. All the products were collected by VAHTS DNA Clean Beads (Vazyme). Then, the end of linearized products was repaired by T4 DNA polymerase (Thermo Fisher Scientific) with 1 mM dNTP mix (Sangon). And the dATP oligo was added at the 3’ end by Dreamtaq polymerase (Thermo Fisher Scientific) with 1 mM dATP (Sangon) to create the 3’ dATP overhang. The products were ligated with adapters which contains 3’ dTTP overhang by fast T4 DNA ligase (Beyotime). The DNA fragments containing the randomized sequences were enriched via PCR using a primer pairing to the adapter and the other primer pairing to the upstream region of the random sequences. The amplified products were purified by VAHTS DNA Clean Beads (Vazyme) and further amplified by primer containing the adapter sequences for Illumina Novaseq PE150 sequencing. In control groups, the plasmid library was directly amplified by two primers covering the randomized sequence for following process. A list of preferred sequences for R2 protein were summarized in Table S4.

#### Reconstitution of complexes

Deactivated R2 protein (D628Y, D996A) was purified as described above. All the complexes were reconstituted by incubating deactivated R2 protein and RNA at RT for 30 minutes in reconstitution buffer (10 mM HEPES-Na pH7.5, 400 mM NaCl, 5 mM MgCl_2_, 2mM DTT). The DNA substrate was added and incubated with RNP at RT for 30 minutes. Then, the assembled sample was purified by size exclusion column (Superose 6 Increase 3.2/300, GE Healthcare) in reconstitution buffer. The peak fraction was stocked at -80 °C after flash-frozen in liquid nitrogen.

#### Cryo-EM sample preparation and data collection

For cryo-EM sample preparation, 4 μL purified complex was loaded onto the graphene oxide grid (Quantifoil Au 1.2/1.3, 400 mesh), which was glow-discharged (HARRICK PLASMA) for 20 seconds at low level after 2 minutes evacuation. The grid was then blotted by a pair of 55 mm filter papers (Ted Pella) at 8 °C with 100% humidity, and flash-frozen in liquid ethane using FEI Vitrobot Marke IV. Cryo-EM data were collected on a Titan Krios electron microscope operated at 300 kV equipped with a Cs-corrector and Gatan K3 direct electron detector with Gatan Quantum energy filter. Micrographs were recorded in counting mode at a nominal magnification of 64,000 x, resulting in a physical pixel size of 1.0979 Å per pixel. The defocus was set between -1.5 μm to -2.5 μm. The total exposure time of each movie stack led to a total accumulated dose of 50 electrons per Å^2^ which fractionated into 32 frames. More parameters for data collection are shown in Table S5.

#### Image processing and 3D reconstruction

The raw dose-fractionated image stacks were 2x Fourier binned, dose-weighted and summed using MotionCor2 (Zheng et al., 2017). The contrast transfer function (CTF) was corrected and bad micrographs were removed manually. The CTF-estimation, blob particles picking, 2D reference-free classification, initial model generation, final 3D refinement and local resolution estimation were performed in cryoSPARC (Punjani et al., 2017). All the 3D reference-based classification were performed in RELION (Kimanius et al., 2016). More details related to data processing were summarized in Figures S2, S5, S8, and Table S5.

#### Model building and refinement

The initial protein model was generated using AlphaFold2 and manually revised in UCSF-Chimera and Coot (Emsley and Cowtan, 2004; Jumper et al., 2021; Pettersen et al., 2004). The initial RNA model was based on the icSHAPE and reported research results, and manually revised based on the cryo-EM density in Coot. The DNA substrates was manually built in Coot based on the cryo-EM density. The knowledge derived from biochemical and sequencing analysis were taking into consideration in the model building and sequence assignment. The complete model was refined against the EM map by PHENIX in real space with secondary structure and geometry restraints (Adams et al., 2010). The final model was validated in PHENIX software package. The structural validation details for the final model are summarized in Table S5.

#### *In vitro* phase separation assays

R2 DrrB protein was purified as described above. The purified DrrB protein were incubated on the ice with 200 ng/mL Alexa Fluor 488 (Thermo Fisher Scientific) for 1 hour in the dark. The fluorescence-conjugated proteins were subsequently diluted with dilution buffer (75 to 300 mM NaCl, 25 mM tris-HCl 7.5) to reach the final concentration of 0.5 to 10 μM. The protein phase separation at different protein concentration and salt concentration reacted at RT for 20 minutes. And the images of phase-separated droplets were taken using the Olympus IXplore Spin Spinning Disk Microscope.

#### RNA secondary structure determination by icSHAPE

RNA secondary structures were determined as previously described with some updates (Piao et al., 2022). RNA was prepared as in the cryo-EM studies in three conditions: apo RNA, RNA-protein binary complex, and RNA-DNA-protein ternary complex. There were three RNAs: 3’-RNA, 5’-RNA and L-RNA. Reverse transcription primers were designed for these nine samples with specific barcodes. RNA modification reagent NAI-N3 was added to these nine samples to a final concentration 100 mM independently, mixed gently, and incubated at 37 °C for 10 minutes. 35 μL of buffer RLT was added to stop the reaction on ice. The RNA was purified with a RNeasy mini column and eluted with 50 μL of pure water twice. The modified RNA was incubated with DIBO-biotin at 37 °C for 2 hours and then purified with the Zymo RNA kit. Reverse transcription was performed for each sample with its specific primers. Then, all the samples were merged after RNA-cDNA hybrid purification. It was then subjected to RNase I treatment and biotin enrichment for the modified RNA-cDNA hybrid. After RNA degradation, linker ligation, and the second DNA chain synthesis, libraries were amplified using qPCR with about 14-16 cycles. The library was then sequenced using Illumina Novaseq PE 150 sequencing. The sequencing data was split into 9 RNA types according to their specific barcode. Only the left reads were used for the following analysis. The icSHAPE score was calculated according to icSHAPE-pipe methods (Li et al., 2020). The icSHAPE profile was shown by Integrative Genomics Viewer (IGV) tools (Robinson et al., 2011).

