## supplemental for "Structural RNA components supervise the sequential DNA cleavage in R2 retrotransposon"


**List:**

Figure S1-S10

Table S1-S5

Video S1-S2

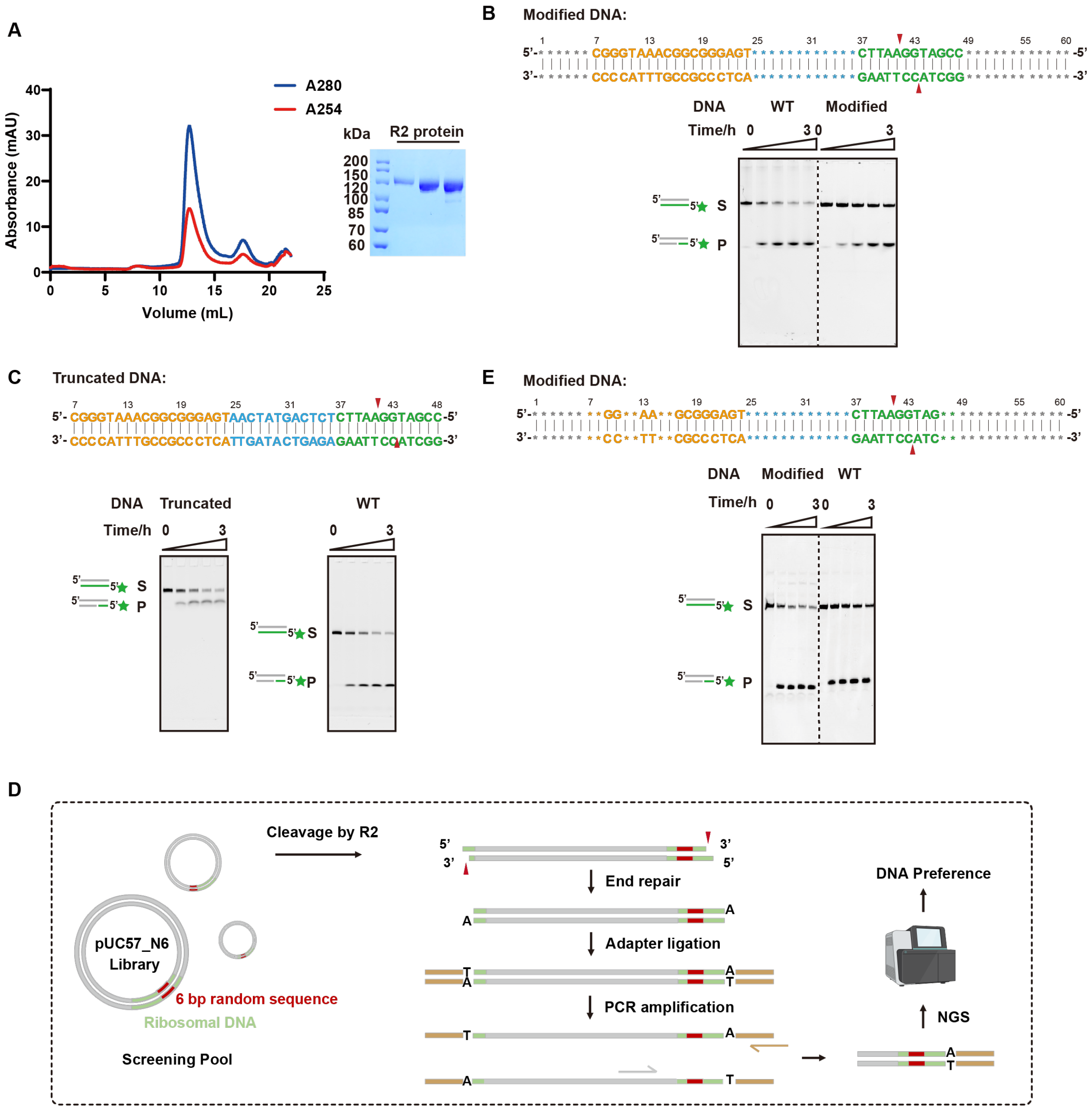


**Figure S1. Biochemical analysis of target recognition by R2 protein. Related to Figure 1.**

(**A**) Purification of R2 protein. Left, the size-exclusion chromatography curves are aligned referring to the injection volume. Right, the peak fractions are analyzed by SDS-PAGE.

(**B**) First-strand cleavage assay by R2 protein for preserving the Dcr and Drr but replacing the sequence of other regions. Top, the sequence of modified DNA. The asterisk indicates the modified nucleotides. The red triangle indicates the cleavage site. Bottom, the denaturing PAGE gel.

(**C**) First-strand cleavage assay by R2 protein for truncating nucleotides in redundant region. Top, the sequence of truncated DNA. The red triangle indicates the cleavage site. Bottom, the denaturing PAGE gel.

(**D**) The deep sequencing workflow for cleavage screening. The pUC57_N6 library contains a 6 bp randomized window, which is cleaved and linearized by R2 protein. The linearized product is repaired to create a dATP overhang at the 3' end. The adapter is ligated through the base pairing between dATP overhang on the linearized products and dTTP overhang on the adapter. PCR is performed to amplify the ligated products and add barcodes for deep sequencing.

(**E**) First-strand cleavage assay by R2 protein for retaining the essential bases but mutating all other ones. Top, the sequence of modified DNA. The asterisk indicates modified nucleotides. The red triangle indicates the cleavage site. Bottom, the denaturing PAGE gel.

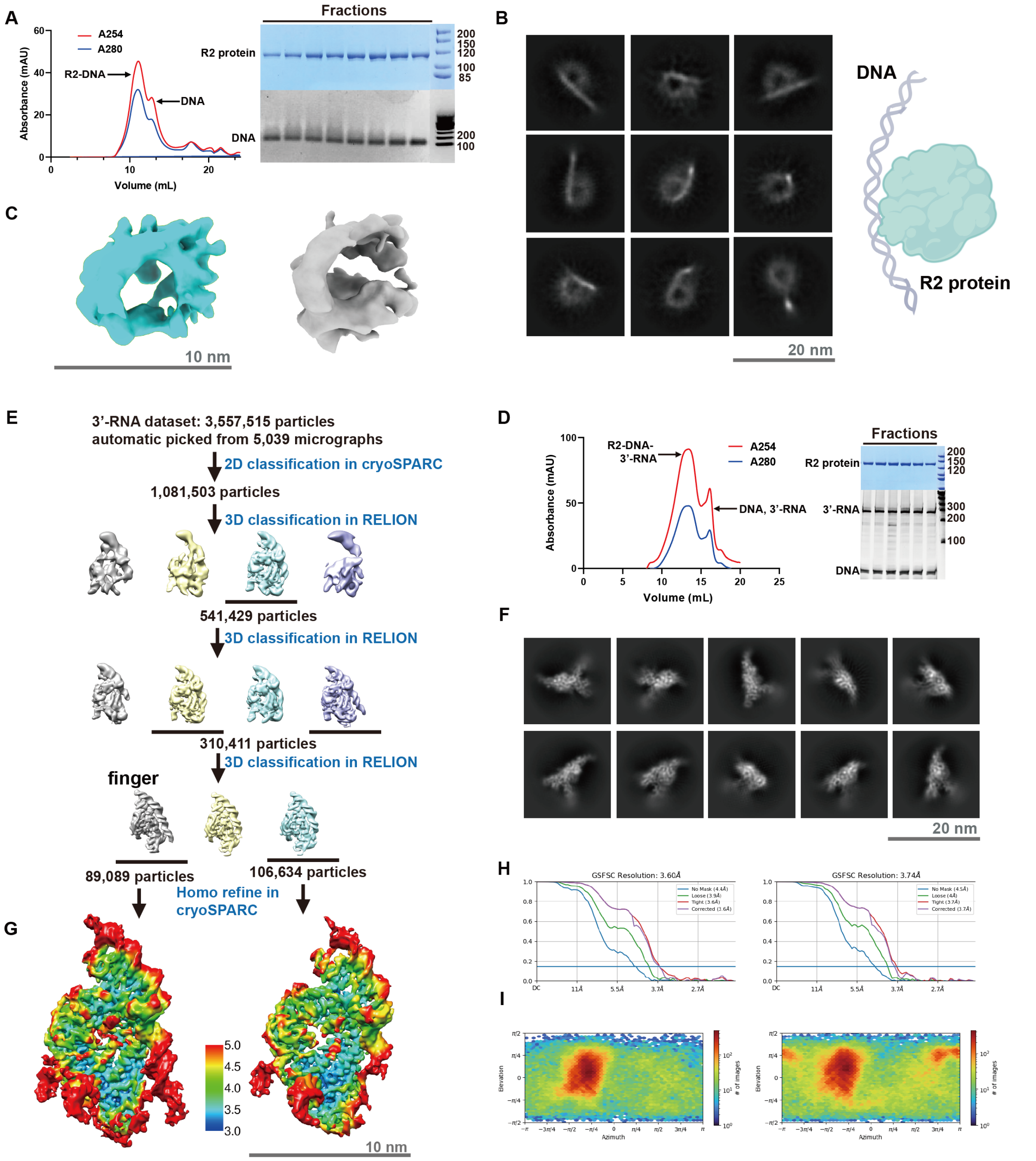


**Figure S2. Cryo-EM analysis of 3'-RNA bound complex. Related to Figure 3.**

(**A**) Reconstitution and purification of R2-DNA binary complex by size-exclusion chromatography. The reconstituted complex is analyzed by SDS-PAGE (right, top) and denaturing PAGE (right, bottom). The peak fractions are used for cryo-EM study.

(**B**) The 2D class-averages of R2-DNA binary complex.

(**C**) The 3D map of R2-DNA binary complex.

(**D**) Reconstitution and purification of 3'-RNA bound complex by size-exclusion chromatography. The reconstituted complex is analyzed by SDS-PAGE (right, top) and denaturing PAGE (right, bottom). The peak fractions are used for cryo-EM study.

(**E**) The workflow for single particle analysis of 3'-RNA bound complex using cryoSPARC and RELION.

(**F**) The 2D class-averages of R2-DNA-3'-RNA ternary complex.

(**G**) The validation of local resolution of 3'-RNA bound complex. The resolution ranging from 3.0 Å to 5.0 Å was shown in the map. Left, the pre-cleavage state. Right, the first-strand cleavage state.

(**H**) Fourier shell correlation (FSC) curves calculated using two independent half-maps. The resolution for the B-factor corrected final maps were 3.6 Å and 3.7 Å, respectively.

(**I**) Euler angle distribution of the refined particles for the final maps of 3'-RNA bound complex. Panel H and I were directly adopted from the standard outputs of cryoSPARC.

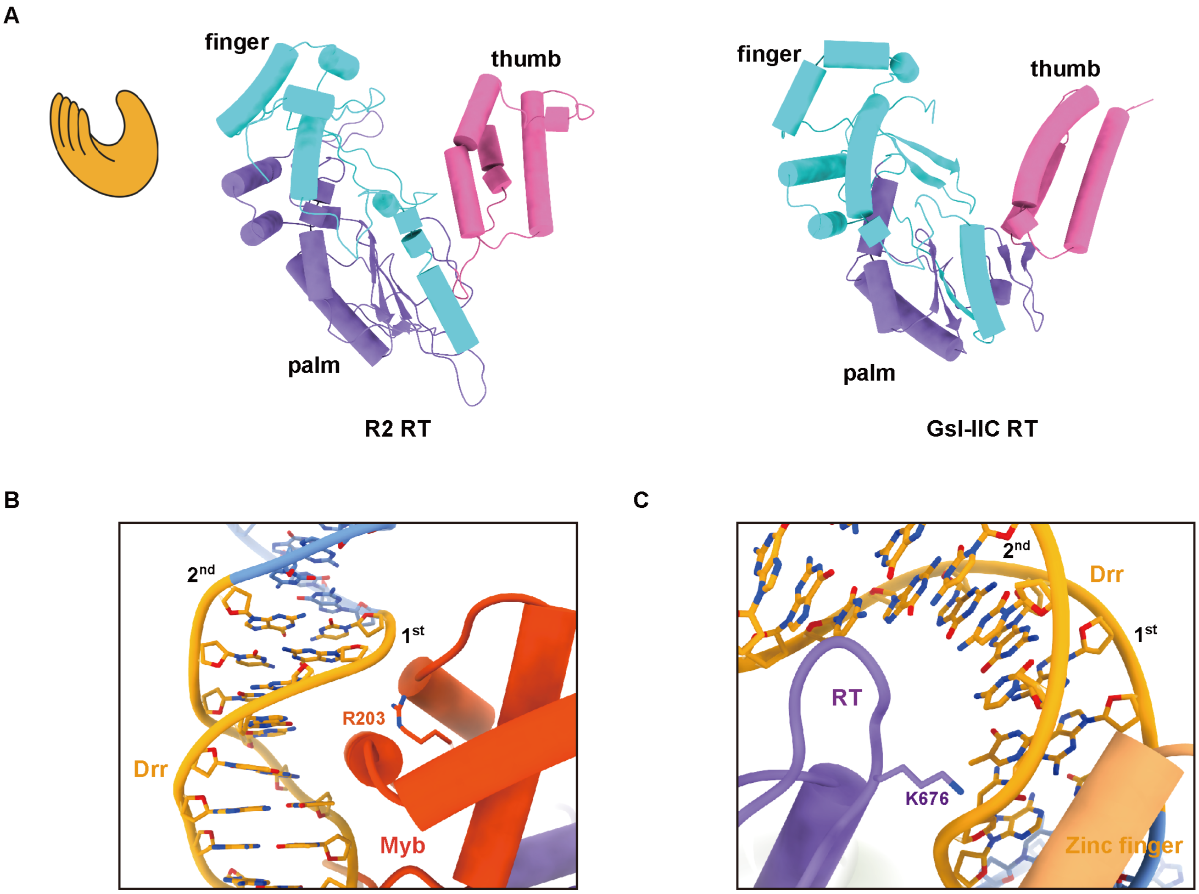


**Figure S3. Structural of 3'-RNA bound complex. Related to Figure 3.**

(**A**) The architecture of RT domain with a right-hand shape. Left, the cartoon model of classic right-hand architecture. Middle, the atomic model of R2 RT. Right, the atomic model of GsI-IIC RT (PDB: 6AR1). The finger is colored blue. The palm is colored purple. The thumb is colored pink.

(**B** and **C**) The structure details for the interaction between DrrB domain and Drr on DNA.

(**B**) The interaction between Myb motif in DrrB domain and the first strand DNA.

(**C**) The interaction between Zinc finger in DrrB domain and the second strand DNA. The residues involving for recognition are labeled.

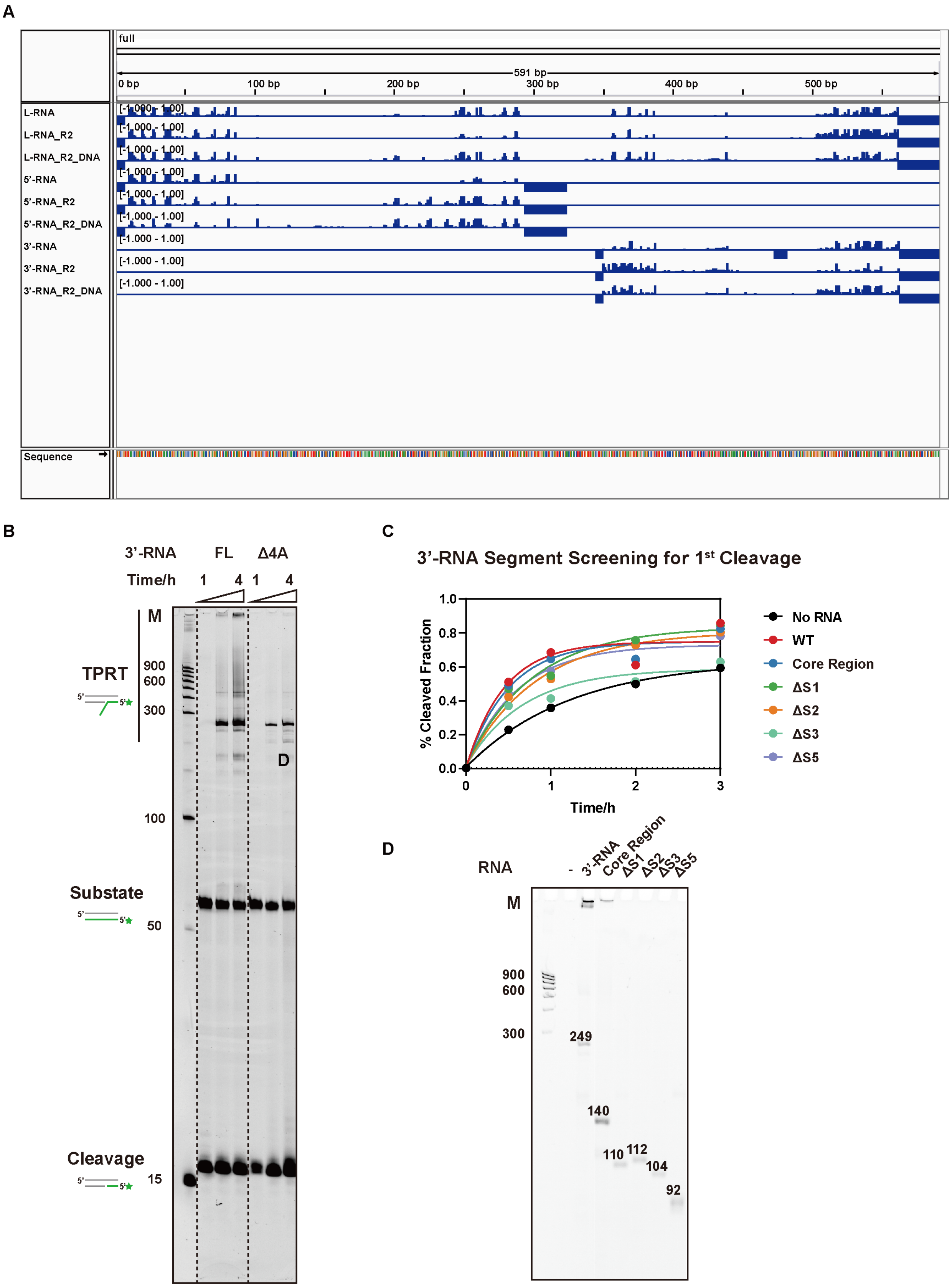

A
full
591 bp
0 bp
100 bp
200 bp
300 bp
400 bp
500 bp
L-RNA
L-RNA_R2
L-RNA_R2_DNA
5'-RNA
5'-RNA_R2
5'-RNA_R2_DNA
3'-RNA
3'-RNA_R2
3'-RNA_R2_DNA
[-1.000 - 1.00]
Sequence
B
3'-RNA
FL
Δ4A
Time/h
1
4
1
4
M
TPRT
900
600
300
D
100
Substate
50
Cleavage
15
C
3'-RNA Segment Screening for 1st Cleavage
% Cleaved Fraction
Time/h
No RNA
WT
Core Region
ΔS1
ΔS2
ΔS3
ΔS5
D
RNA
-
3'-RNA
Core Region
ΔS1
ΔS2
ΔS3
ΔS5
M
900
600
300
249
140
110
112
104
92

**Figure S4. Biochemical analysis of 3'-RNA. Related to Figure 4.**

(**A**) RNA secondary structure analysis via icSHAPE. The data are scaled from 0 (no reactivity) and 1 (maximum reactivity). The reactivity of L-RNA (591 nt) is drawn from 0 to 591. The reactivity of 5'-RNA (323 nt) is drawn from 0 to 323. The reactivity of 3'-RNA (249 nt) is drawn from 343 to 591.

(**B**) The TPRT assay for shortening the length of 3'-RNA polyA tail. FL, full length 3'-RNA (249 nt). Δ4A, 3'-RNA deleting A245 to A249 at the 3'-end (245 nt).

(**C**) The plot of cleavage efficiency for the first-strand cleavage assay in Figure 4E.

(**D**) The RNA staining of the denaturing PAGE gel for the TPRT assay in Figure 4F.

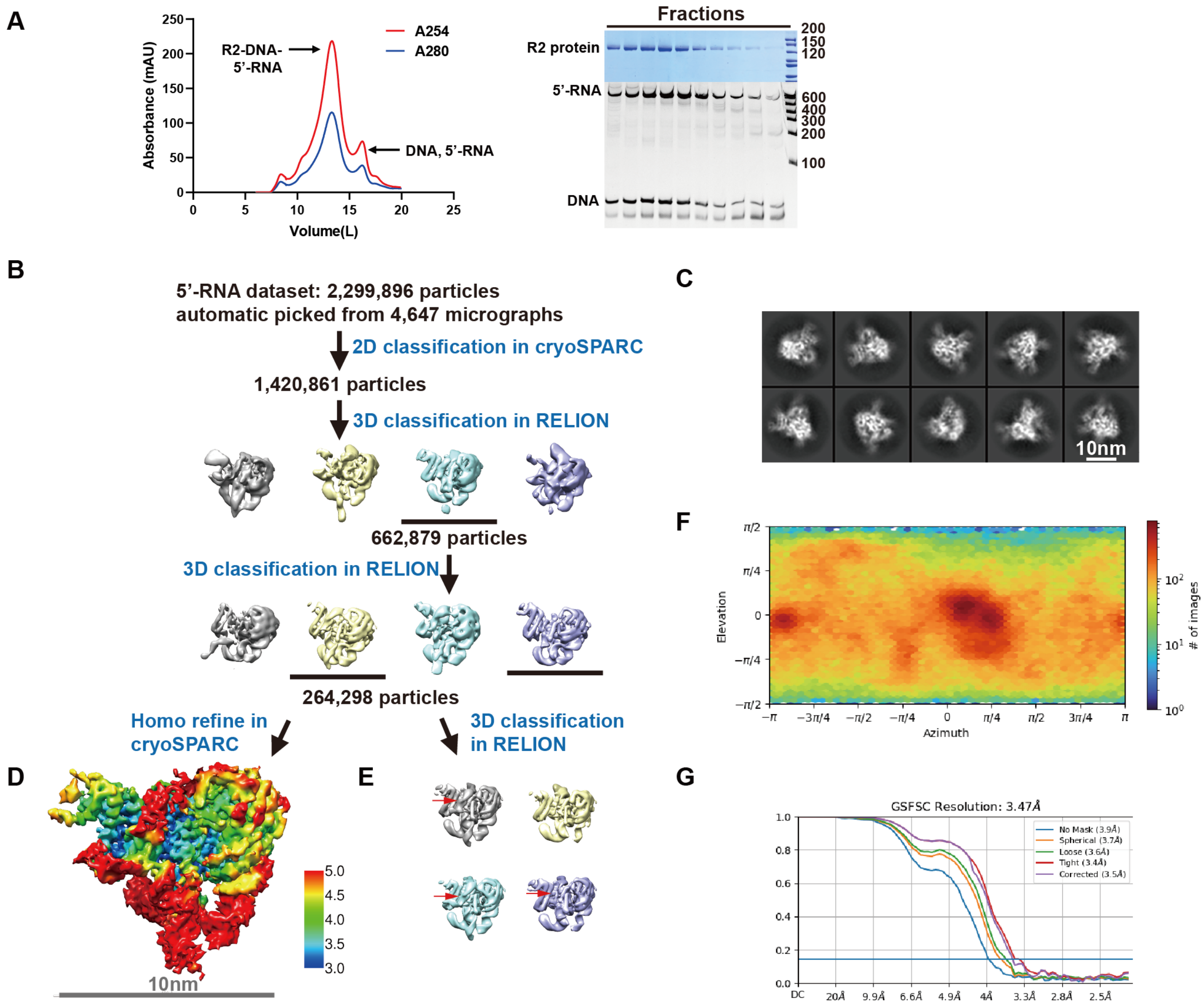


**Figure S5. Cryo-EM analysis of 5'-RNA bound complex. Related to Figure 5.**

(**A**) Reconstitution and purification of 5'-RNA bound complex by size-exclusion chromatography. The reconstituted complex is analyzed by SDS-PAGE (right, top) and denaturing PAGE (right, bottom). The peak fractions are used for cryo-EM study.

(**B**) The workflow for single particle analysis of 5'-RNA bound complex using cryoSPARC and RELION.

(**C**) The 2D-averages of 5'-RNA bound complex.

(**D**) The validation of local resolution of R2-DNA-5'-RNA ternary complex. The resolution ranging from 3.0 Å to 5.0 Å was shown in the map.

(**E**) The 3D-classification of 5'-RNA bound complex. The red arrow indicates the different density in the map.

(**F**) Euler angle distribution of the refined particles for the final map of 5'-RNA bound complex. Panel F and G were directly adopted from the standard outputs of cryoSPARC.

(**G**) Fourier shell correlation (FSC) curves calculated using two independent half-maps. The resolution for the B-factor corrected final map was 3.5 Å.

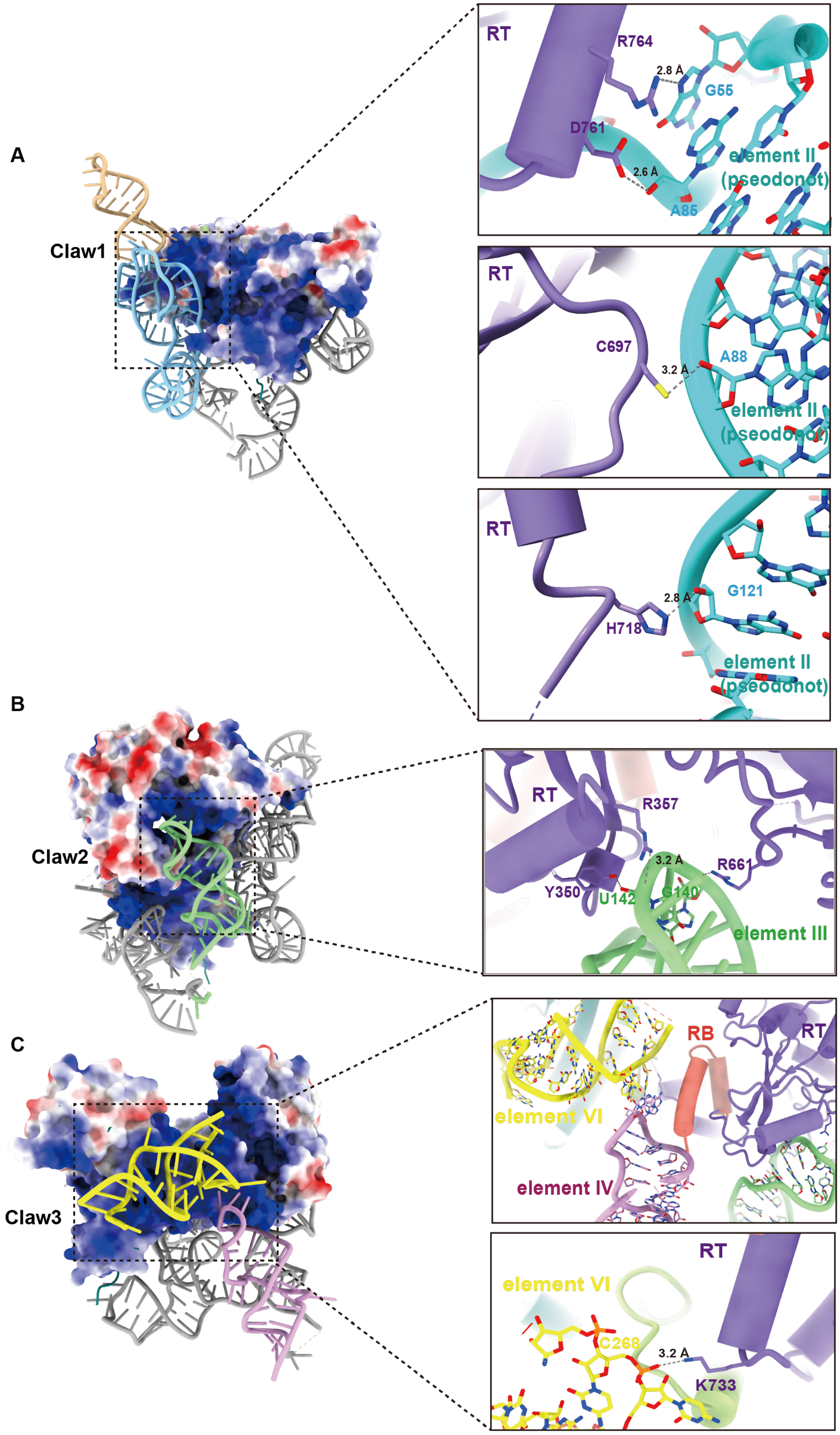
A
Claw1
RT
R764
2.8 Å
G55
D761
2.6 Å
A85
element II
(pseodonot)
RT
C697
3.2 Å
A88
element II
(pseodonot)
RT
2.8 Å
G121
H718
element II
(pseodonot)
B
Claw2
RT
R357
3.2 Å
R661
Y350
U142
G140
element III
C
Claw3
RB
RT
element VI
element IV
RT
element VI
C268
3.2 Å
K733

**Figure S6. The three-claw architecture of 5'-RNA. Related to Figure 5.**

(**A-C**) Left, the electrostatic interaction interface between protein and 5'-RNA. Blue indicates a positive charge while red indicates a negative charge. Right, the structure details for the interaction between the claws and R2 protein. The hydrogen-bonds interaction is shown by dashed lines and labeled with atom distance. The protein domains and 5'-RNA elements are colored corresponding to Figure 5.

(**A**) The architecture and interactions for claw1 of 5'-RNA.

(**B**) The architecture and interactions for claw2 of 5'-RNA.

(**C**) The architecture and interactions for claw3 of 5'-RNA.

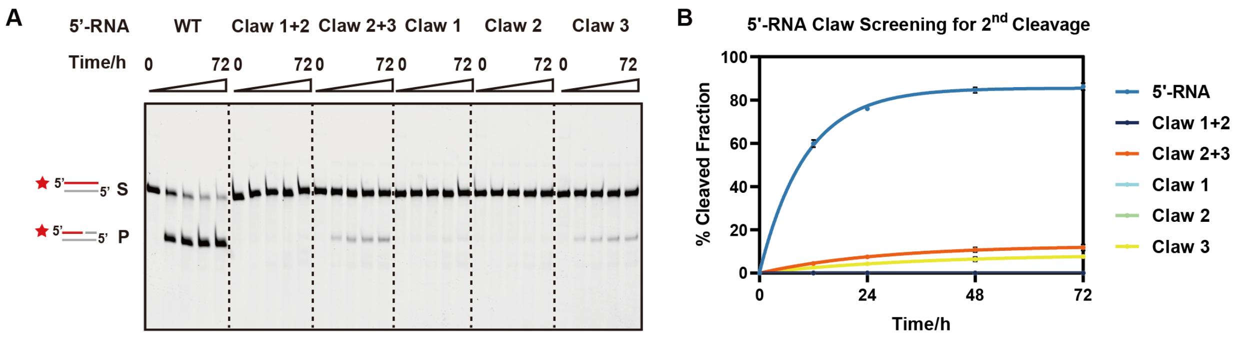


**Figure S7. The biochemical analysis of 5'-RNA claws. Related to Figure 5.**

(**A**) Second-strand cleavage for truncated 5'-RNA.

(**B**) The plot of cleavage efficiency for the second-strand cleavage assay in panel A. (n=3 each; means ± SD) (S, substrates. P, products. Red colored pentagram indicates FAM labeling at 5' end of DNA.)

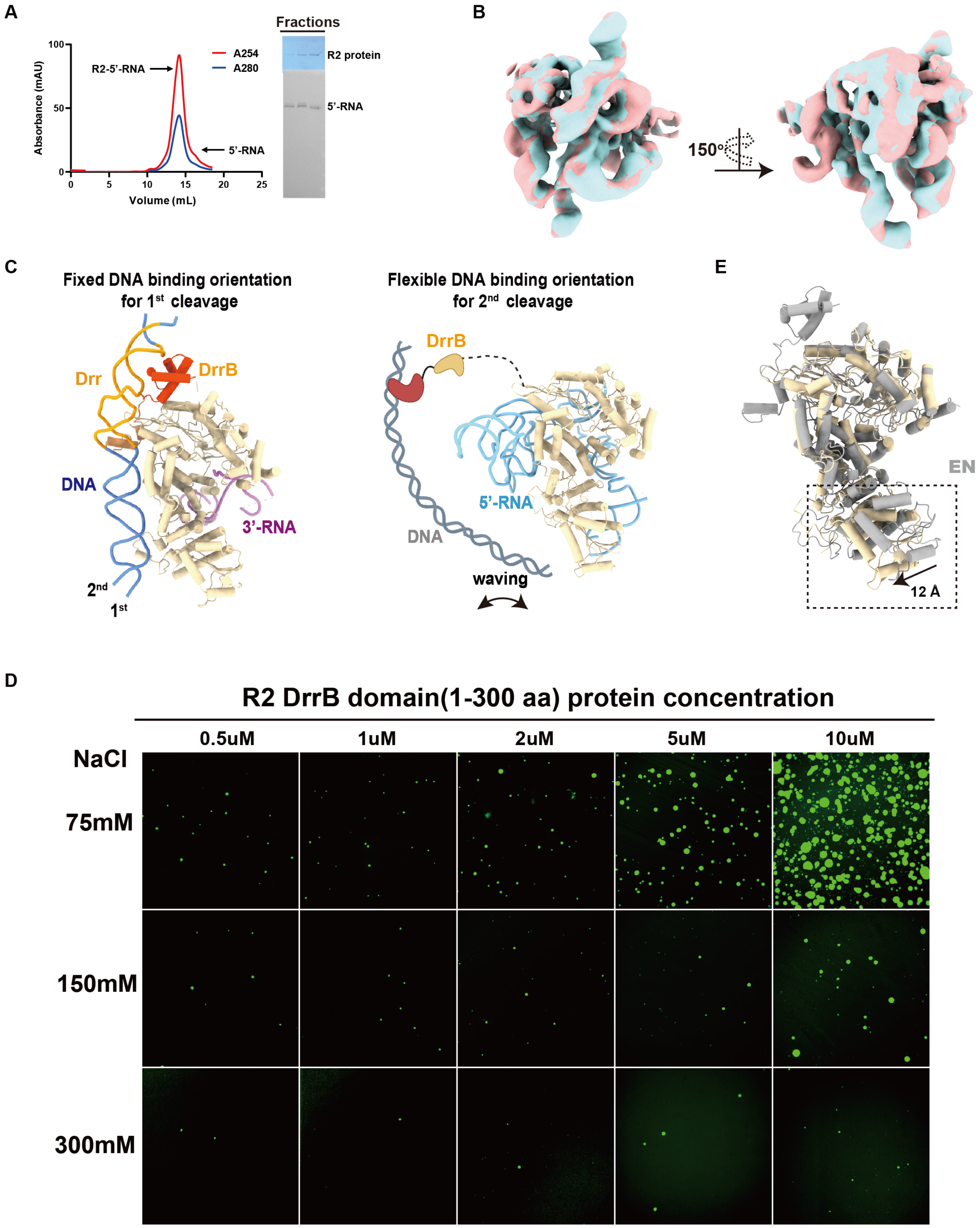


**Figure S8. Structural and functional analysis of DrrB domain. Related to Figure 5.**

(**A**) Reconstitution and purification of R2-5'-RNA binary complex by size-exclusion chromatography. The reconstituted complex is analyzed by SDS-PAGE (right, top) and denaturing PAGE (right, bottom). The peak fractions are used for cryo-EM study.

(**B**) The cryo-EM map alignment of R2-5'-RNA binary complex and R2-DNA-5'-RNA ternary complex. The binary complex is colored blue. The ternary complex is colored pink.

(**C**) The comparison of 3'-RNA bound complex and 5'-RNA bound complex. Left, the atomic model for 3'-RNA bound complex. Right, the atomic model for 5'-RNA bound complex. The cartoon model indicates the hypothetical position of the DNA and DrrB domain, which are not observed in the 3D structure.

(**D**) *In vitro* phase separation assay for DrrB domain. The concentration of DrrB is ranged from 0.5 to 10 uM. The concentration of NaCl is ranged from 75 to 300 mM.

(**E**) The alignment of protein in 3'-RNA bound complex and protein in 5'-RNA bound complex. Protein in 3'-RNA bound complex is colored gray. Protein in 5'-RNA ternary complex is colored wheat.

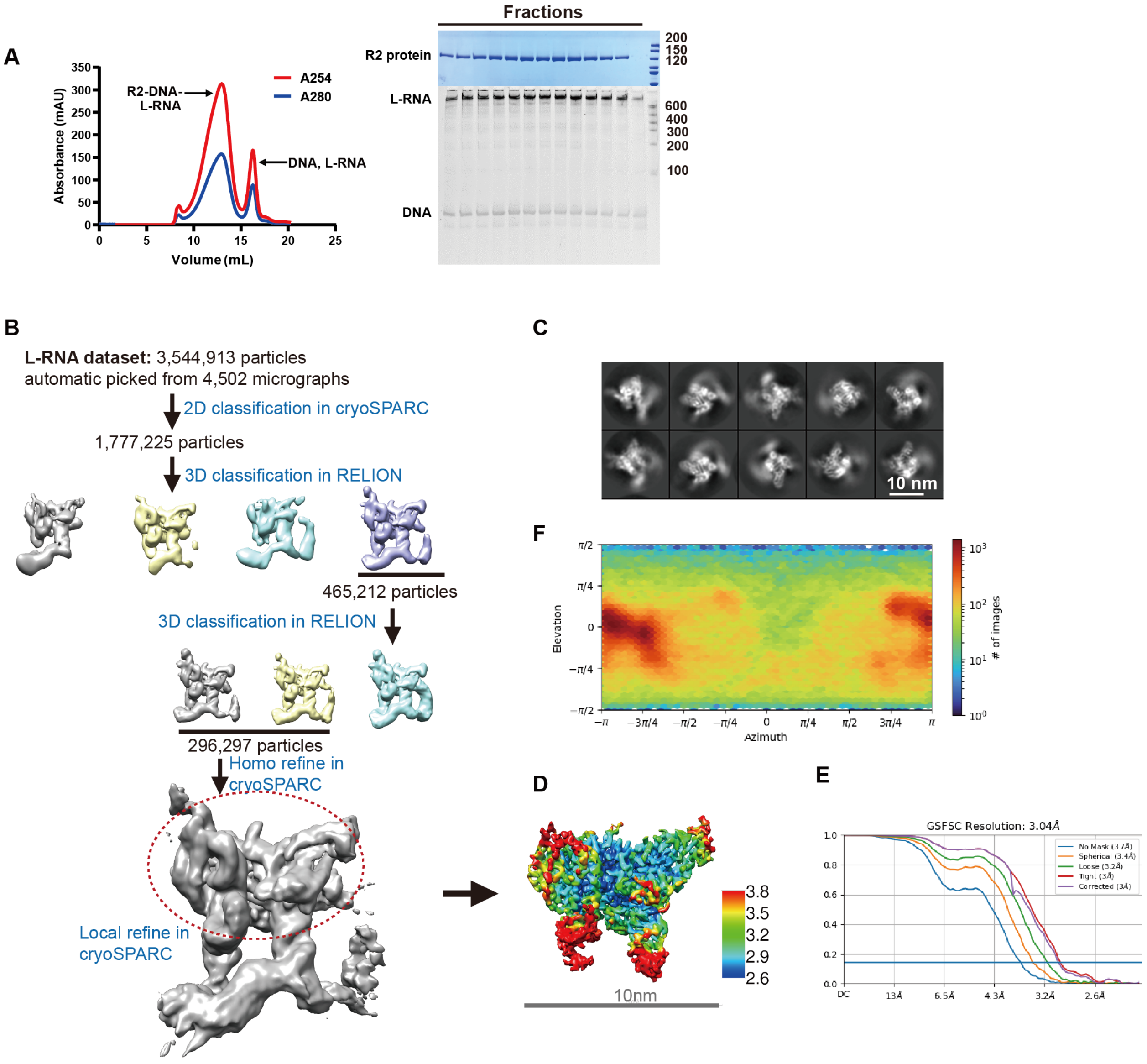


**Figure S9. Cryo-EM analysis of L-RNA bound complex. Related to Figure 6.**

(**A**) Reconstitution and purification of L-RNA bound complex by size-exclusion chromatography. The reconstituted complex is analyzed by SDS-PAGE (right, top) and denaturing PAGE (right, bottom). The peak fractions are used for cryo-EM study.

(**B**) The workflow for single particle analysis of L-RNA bound complex using cryoSPARC and RELION.

(**C**) The representative 2D class-averages of L-RNA complex.

(**D**) The validation of local resolution of L-RNA bound complex. The resolution ranging from 2.6 Å to 3.8 Å is shown in the map.

(**E**) Fourier shell correlation (FSC) curves calculated using two independent half-maps. The resolution for the B-factor corrected final map is 3.0 Å.

(**F**) Euler angle distribution of the refined particles for the final map of L-RNA bound complex. Panel E and F are directly adopted from the standard outputs of cryoSPARC.

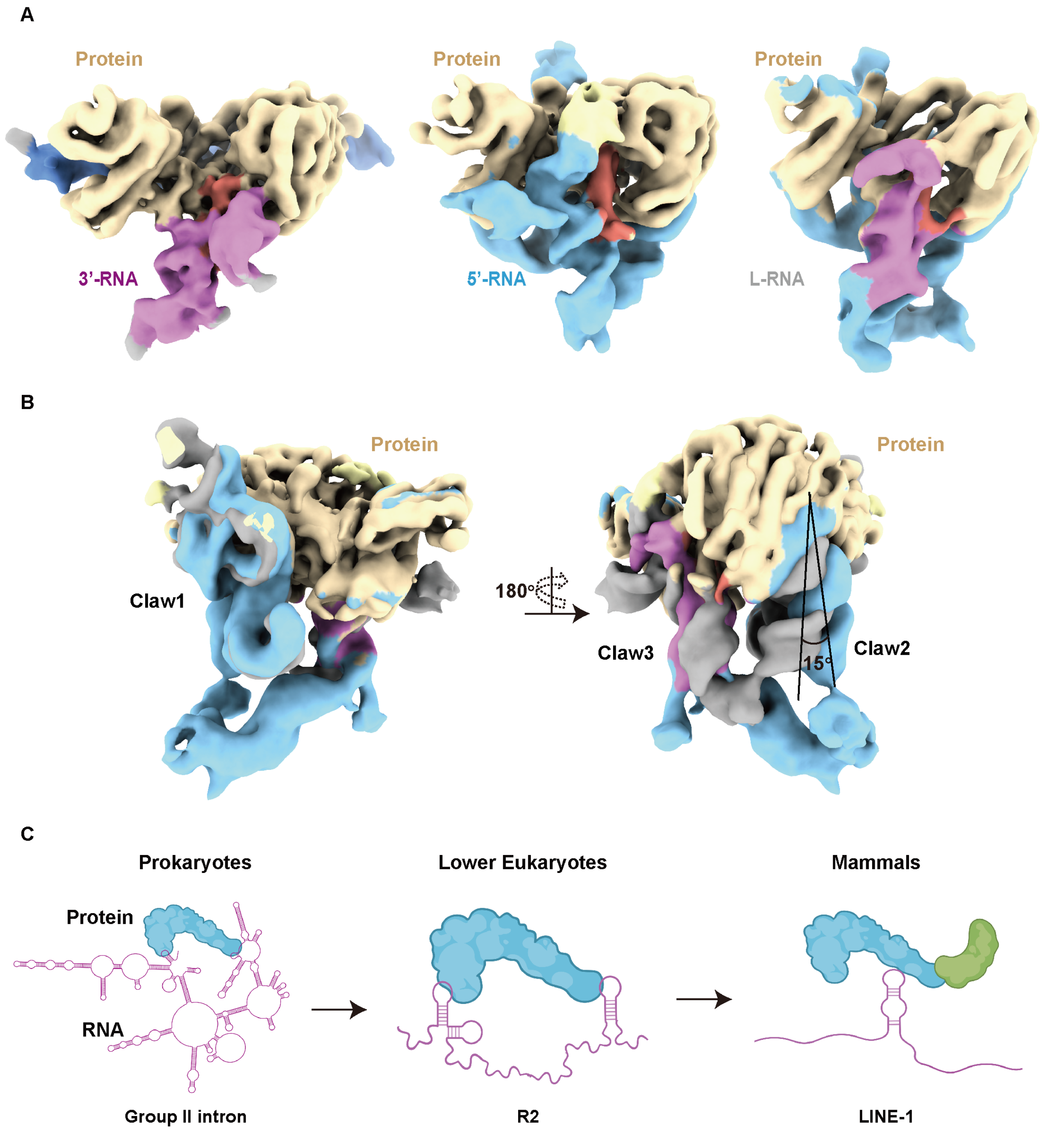


**Figure S10. Structural comparison of 3'-RNA, 5'-RNA and L-RNA bound complex. Related to Figure 6.**

(**A**) The 3D map comparison of proteins and RNA in three bound complexes. The protein is colored wheat and the RB domain is colored red. The 3'-RNA is colored pink. The 5'-RNA is colored blue.

(**B**) The structure alignment of L-RNA and 5’-RNA bound complex. The protein in L-RNA and 5’-RNA bound complex is colored wheat. The 3’-RNA density observed in L-RNA bound complex is colored pink, the 5’-RNA density in L-RNA is colored blue and the 5’-RNA density in 5’-RNA bound complex is colored gray.

(**C**) The hypothetical trend for co-evolution between RNA and protein from prokaryotes, lower eukaryotes to mammals.

**Table S1. Plasmids generated in this study.**

| ID | Assay | Features | Selection marker |
|---|---|---|---|
| p-001 | Protein purification | pET28a-backbone derived plasmids containing N-terminal 6xHis-SUMO tagged R2Bm in MCS. | Kanamycin |
| p-002 | Protein purification | pET28a-backbone derived plasmids containing N-terminal 6xHis-SUMO tagged R2Bm(D628Y D996A) in MCS. | Kanamycin |
| p-003 | Protein purification | pET28a-backbone derived plasmids containing N-terminal 6xHis-SUMO tagged DrrB domain in MCS. | Kanamycin |
| p-004 | *In vitro* plasmids cleavage screening | pUC57-backbone derived plasmids containing native ribosomal DNA and randomized window CAAGCGNNNNNNAACGGCGGGAGTAACTATGACTCTCTTAAGGTAGCCAAATGCCTCGTC. | Ampicillin |
| p-005 | *In vitro* plasmids cleavage screening | pUC57-backbone derived plasmids containing native ribosomal DNA and randomized window CAAGCGCGGGTANNNNNNGGGAGTAACTATGACTCTCTTAAGGTAGCCAAATGCCTCGTC. | Ampicillin |
| p-006 | *In vitro* plasmids cleavage screening | pUC57-backbone derived plasmids containing native ribosomal DNA and randomized window CAAGCGCGGGTAAACGGCNNNNNNAACTATGACTCTCTTAAGGTAGCCAAATGCCTCGTC. | Ampicillin |
| p-007 | *In vitro* plasmids cleavage screening | pUC57-backbone derived plasmids containing native ribosomal DNA and randomized window CAAGCGCGGGTAAACGGCGGGAGTAACTATNNNNNNCTTAAGGTAGCCAAATGCCTCGTC. | Ampicillin |
| p-009 | *In vitro* plasmids cleavage screening | pUC57-backbone derived plasmids containing native ribosomal DNA and randomized window CAAGCGCGGGTAAACGGCGGGAGTAACTATGACTCTNNNNNNGTAGCCAAATGCCTCGTC. | Ampicillin |
| p-011 | *In vitro* plasmids cleavage screening | pUC57-backbone derived plasmids containing native ribosomal DNA and randomized window CAAGCGCGGGTAAACGGCGGGAGTAACTATGACTCTCTTAAGNNNNNNAAATGCCTCGTC. | Ampicillin |

**Table S2. RNA sequences used in this study**

| ID # | Description | Sequences (5'-3') |
|---|---|---|
| RNA-001 | 3'-RNA sequence for biochemical assay and cryo-EM analysis | GGCCUUGCACAGUAGUCCAGCGGUAAGGGUGUAGAUC AGGCCCGUCUGUUUCUCCCCCGGAGCUCGCUCCCUUG GCUUCCCUUAUAUAUUUUAACAUCAGAAACAGACAUU AAACAUCUACUGAUCCAAUUUCGCCGGCGUACGGCCA CGAUCGGGAGGGUGGGAAUCUCGGGGGUCUUCCGAUC CUAAUCCAUGAUGAUUACGACCUGAGUCACUAAAGAC GAUGGCAUGAUGAUCCGGCGAUGAAAA |
| RNA-002 | 5'-RNA sequence for biochemical assay and cryo-EM analysis | GGGCCGGUGUAACCCGGAUGGCUGUACACGUGGUAAA CACGUGACAGCAGCCCCGAUGGACGGACCGCGAGGAC CGUCAAGCCUAGCAGGUACCUUCGGGUGGGGCCUUGC GAUACCUGCGGGCGAACCCUGUGGUCGGGUUUGCAGC CCGGCCACAGUGGGUUUUUUUCCUGUUGCAAAAAAGU CAAAUAAAGAAAAUAGACCUGAAGCCUCUGGCCUCCC GCUGGAGUCAGAGAGGACAGGCGAUAACCCGACUGUG CGGGGUUCCGCCGGCGCAGAUCCUGUGGGUCAGGAUG CGCCUGGUUGGACCUGCCAGUUCUGCG |
| RNA-003 | L-RNA sequence for biochemical assay and cryo-EM analysis | GGGCCGGUGUAACCCGGAUGGCUGUACACGUGGUAAA CACGUGACAGCAGCCCCGAUGGACGGACCGCGAGGAC CGUCAAGCCUAGCAGGUACCUUCGGGUGGGGCCUUGC GAUACCUGCGGGCGAACCCUGUGGUCGGGUUUGCAGC CCGGCCACAGUGGGUUUUUUUCCUGUUGCAAAAAAGU CAAAUAAAGAAAAUAGACCUGAAGCCUCUGGCCUCCC GCUGGAGUCAGAGAGGACAGGCGAUAACCCGACUGUG CGGGGUUCCGCCGGCGCAGAUCCUGUGGGUCAGGAUG CGCCUGGUUGGACCUGCCAGUUCUGCGAACGAACCUU CGUUGGUUGAGCCUUGCACAGUAGUCCAGCGGUAAGG GUGUAGAUCAGGCCCGUCUGUUUCUCCCCCGGAGCUC GCUCCCUUGGCUUCCCUUAUAUAUUUUAACAUCAGAA ACAGACAUUAAACAUCUACUGAUCCAAUUUCGCCGGC GUACGGCCACGAUCGGGAGGGUGGGAAUCUCGGGGGU CUUCCGAUCCUAAUCCAUGAUGAUUACGACCUGAGUC ACUAAAGACGAUGGCAUGAUGAUCCGGCGAUGAAAA |
| RNA-004 | Core Region of 3'-RNA sequence for biochemical assay | GGCCUUGCACAGUAGUCCAGCGGUAAGGGUGUAGAUC AGGCCCGUCUGUUUCUAACAUCAGAAACAGACAUUAA ACAUCUACUGAUCCAAUUUCGCCGGCGUACGGCCAAG ACGAUGGCAUGAUGAUCCGGCGAUGAAAA |
| RNA-005 | ΔS1 of 3'-RNA sequence for biochemical assay | GUAGAUCAGGCCCGUCUGUUUCUAACAUCAGAAACAG ACAUUAAACAUCUACUGAUCCAAUUUCGCCGGCGUAC GGCCAAGACGAUGGCAUGAUGAUCCGGCGAUGAAAA |
| RNA-006 | ΔS2 of 3'-RNA sequence for biochemical assay | GGCCUUGCACAGUAGUCCAGCGGUAAGGGUGUAGUCU GUUUCUAACAUCAGAAACAGACCAAUUUCGCCGGCGU ACGGCCAAGACGAUGGCAUGAUGAUCCGGCGAUGAAA A |
| RNA-007 | ΔS3 of 3'-RNA sequence for biochemical assay | GGCCUUGCACAGUAGUCCAGCGGUAAGGGUGUAGAUC AGGCCCUACUGAUCCAAUUUCGCCGGCGUACGGCCAA GACGAUGGCAUGAUGAUCCGGCGAUGAAAA |
| RNA-008 | ΔS5 of 3'-RNA sequence for biochemical assay | GGCCUUGCACAGUAGUCCAGCGGUAAGGGUGUAGAUC AGGCCCGUCUGUUUCUAACAUCAGAAACAGACAUUAA ACAUCUACUGAUCCAAUU |
| RNA-009 | Δ4A of 3'-RNA sequence for biochemical assay | GGCCUUGCACAGUAGUCCAGCGGUAAGGGUGUAGAUC AGGCCCGUCUGUUUCUCCCCCGGAGCUCGCUCCCUUG GCUUCCCUUAUAUAUUUUAACAUCAGAAACAGACAUU AAACAUCUACUGAUCCAAUUUCGCCGGCGUACGGCCA |

| | | |
|---|---|---|
| | | CGAUCGGGAGGGUGGGAAUCUCGGGGGUCUUCCGAUC CUAAUCCAUGAUGAUUACGACCUGAGUCACUAAAGAC GAUGGCAUGAUGAUCCGGCGAUG |
| RNA-010 | Claw 1+2 of 5’-RNA sequence for biochemical analysis | GGGCCGGUGUAACCCGGAUGGCUGUACACGUGGUAAA CACGUGACAGCAGCCCCGAUGGACGGACCGCGAGGAC CGUCAAGCCUAGCAGGUACCUUCGGGUGGGGCCUUGC GAUACCUGCGGGCGAACCCUGUGGUCGGGUUUGCAGC CCGGCCACAGUGGGUUU |
| RNA-011 | Claw 2+3 of 5’-RNA sequence for biochemical analysis | GAACCCUGUGGUCGGGUUUGCAGCCCGGCCACAGUGG GUUUUUUUCCUGUUGCAAAAAAGUCAAAUAAAGAAA AUAGACCUGAAGCCUCUGGCCUCCCGCUGGAGUCAGA GAGGACAGGCGAUAACCCGACUGUGCGGGGUUCCGCC GGCGCAGAUCCUGUGGGUCAGGAUGCGCCUGGUUGGA CCUGCCAGUUCUGCG |
| RNA-012 | Claw 1 of 5’-RNA sequence for biochemical analysis | GGGCCGGUGUAACCCGGAUGGCUGUACACGUGGUAAA CACGUGACAGCAGCCCCGAUGGACGGACCGCGAGGAC CGUCAAGCCUAGCAGGUACCUUCGGGUGGGGCCUUGC GAUACCUGCGGGC |
| RNA-013 | Claw 2 of 5’-RNA sequence for biochemical analysis | GAACCCUGUGGUCGGGUUUGCAGCCCGGCCACAGUGG GUUU |
| RNA-014 | Claw 3 of 5’-RNA sequence for biochemical analysis | UUUCCUGUUGCAAAAAAGUCAAAUAAAGAAAAUAGA CCUGAAGCCUCUGGCCUCCCGCUGGAGUCAGAGAGGA CAGGCGAUAACCCGACUGUGCGGGGUUCCGCCGGCGC AGAUCCUGUGGGUCAGGAUGCGCCUGGUUGGACCUGC CAGUUCUGCG |

**Table S3. DNA Oligonucleotides and target sequences used in this study.**

| ID # | Assay | Description | Sequences (5'-3') |
|---|---|---|---|
| primer-001 | Cy5-labeled dsDNA cleavage | Reverse PCR primer for cy5-labeled dsDNA target amplification | Cy5-GACACGACGCTTAG |
| primer-002 | Cy5-labeled dsDNA cleavage | Forward PCR primer for cy5-labeled dsDNA target amplification | Cy5-CAAGCGCGGGTAAAC |
| primer-003 | Cy5-labeled dsDNA cleavage for window mutation screening assay | Reverse PCR primes for cy5-labeled dsDNA target amplification | Cy5-CTGTGCTGCGAATC |
| primer-004 | Cy5-labeled dsDNA cleavage for window mutation screening assay | Reverse PCR primes for cy5-labeled dsDNA target amplification | Cy5-GACGAGGCATTTAC |
| primer-005 | Cy5-labeled dsDNA cleavage for window mutation screen ing assay | Reverse PCR primes for cy5-labeled dsDNA target amplification | Cy5-GACGAGAATACCGG |
| primer-006 | Cy5-labeled dsDNA cleavage for window mutation screening assay | Reverse PCR primes for cy5-labeled dsDNA target amplification | Cy5-CGATGCGCATTTGG |
| target-001 | Cy5-labeled dsDNA cleavage | Forward template for Cy5-labeled native dsDNA target amplification and dsDNA annealing | CAAGCGCGGGTAAACGGC GGGAGTAACTATGACTCTC TTAAGGTAGCCAAATGCCT CGTC |
| target-002 | Cy5-labeled dsDNA cleavage for window mutation screening assay | Forward template for Cy5-labeled W1 dsDNA target amplification | GCCTACCGGGTAAACGGC GGGAGTAACTATGACTCTC TTAAGGTAGCCAAATGCCT CGTC |
| target-003 | Cy5-labeled dsDNA cleavage for window mutation screening assay | Forward template for Cy5-labeled W2 dsDNA target amplification | CAAGCGACTCGGAACGGC GGGAGTAACTATGACTCTC TTAAGGTAGCCAAATGCCT CGTC |
| target-004 | Cy5-labeled dsDNA cleavage for window mutation screening assay | Forward template for Cy5-labeled W3 dsDNA target amplification | CAAGCGCGGGTAGTGACG GGGAGTAACTATGACTCTC TTAAGGTAGCCAAATGCCT CGTC |
| target-005 | Cy5-labeled dsDNA cleavage for window mutation screening assay | Forward template for Cy5-labeled W4 dsDNA target amplification | CAAGCGCGGGTAAACGGC CTCGAGAACTATGACTCTC TTAAGGTAGCCAAATGCCT CGTC |
| target-006 | Cy5-labeled dsDNA cleavage for window mutation screening assay | Forward template for Cy5-labeled W5 dsDNA target amplification | CAAGCGCGGGTAAACGGC GGGAGTTTAGTAGACTCTC TTAAGGTAGCCAAATGCCT CGTC |
| target-007 | Cy5-labeled dsDNA cleavage for window mutation screening assay | Forward template for Cy5-labeled W6 dsDNA target amplification | CAAGCGCGGGTAAACGGC GGGAGTAACTATACTCACC TTAAGGTAGCCAAATGCCT CGTC |
| target-008 | Cy5-labeled dsDNA cleavage for window mutation screening assay | Forward template for Cy5-labeled W7 dsDNA target amplification | CAAGCGCGGGTAAACGGC GGGAGTAACTATGACTCTT GAGTTGTAGCCAAATGCCT CGTC |
| target-009 | Cy5-labeled dsDNA cleavage for window mutation screening assay | Forward template for Cy5-labeled W8 dsDNA target amplification | CAAGCGCGGGTAAACGGC GGGAGTAACTATGACTCTC |

| | | | |
|---|---|---|---|
| | | | TTAAGCCCAGTAAATGCCT CGTC |
| target-010 | Cy5-labeled dsDNA cleavage for window mutation screening assay | Forward template for Cy5-labeled W9 dsDNA target amplification | CAAGCGCGGGTAAACGGC GGGAGTAACTATGACTCTC TTAAGGTAGCCGGTATTCT CGTC |
| target-011 | Cy5-labeled dsDNA cleavage for window mutation screening assay | Forward template for Cy5-labeled W10 dsDNA target amplification | CAAGCGCGGGTAAACGGC GGGAGTAACTATGACTCTC TTAAGGTAGCCAAATGCGC ATCG |
| target-012 | Cy5-labeled dsDNA cleavage for modified DNA assay | Forward template for Cy5-labeled Replace1 dsDNA target amplification | GCTCACCGGGTAAACGGC GGGAGTGCAATATCTGTAC TTAAGGTAGGATTCGCAGC ACAG |
| target-013 | Cy5-labeled dsDNA cleavage for spacer length screening assay | Forward template for Cy5-labeled 8bp spacer dsDNA target amplification | CAAGCGCGGGTAAACGGC GGGAGTGCAATATCCTTAA GGTAGCCAAATGCCTCGTC |
| target-014 | Cy5-labeled dsDNA cleavage for spacer length screening assay | Forward template for Cy5-labeled 10 bp spacer dsDNA target amplification | CAAGCGCGGGTAAACGGC GGGAGTGCAATATCTGCTT AAGGTAGCCAAATGCCTC GTC |
| target-015 | Cy5-labeled dsDNA cleavage for spacer length screening assay | Forward template for Cy5-labeled 11 bp spacer dsDNA target amplification | CAAGCGCGGGTAAACGGC GGGAGTGCAATATCTGTCT TAAGGTAGCCAAATGCCTC GTC |
| target-016 | Cy5-labeled dsDNA cleavage for spacer length screening assay | Forward template for Cy5-labeled 12 bp spacer dsDNA target amplification | CAAGCGCGGGTAAACGGC GGGAGTGCAATATCTGTAC TTAAGGTAGCCAAATGCCT CGTC |
| target-017 | Cy5-labeled dsDNA cleavage for spacer length screening assay | Forward template for Cy5-labeled 13 bp spacer dsDNA target amplification | CAAGCGCGGGTAAACGGC GGGAGTGCAATATCTGTA GCTTAAGGTAGCCAAATGC CTCGTC |
| target-018 | Cy5-labeled dsDNA cleavage for spacer length screen assay | Forward template for Cy5-labeled 14 bp spacer dsDNA target amplification | CAAGCGCGGGTAAACGGC GGGAGTGCAATATCTGTA GCCTTAAGGTAGCCAAATG CCTCGTC |
| target-019 | Cy5-labeled dsDNA cleavage for spacer length screen assay | Forward template for Cy5-labeled 16 bp spacer dsDNA target amplification | CAAGCGCGGGTAAACGGC GGGAGTGCAATATCTGTAG CTCCTTAAGGTAGCCAAAT GCCTCGTC |
| target-020 | Cy5-labeled dsDNA cleavage for truncated assay | Cy5-labeled ssDNA for truncated dsDNA annealing | Cy5-GGCTACCTTAAGAGAGTCA TAGTTACTCCCGCCGTTTA CCCG |
| target-021 | Cy5-labeled dsDNA cleavage for truncated assay | Non-labeled ssDNA for truncated dsDNA annealing | CGGGTAAACGGCGGGAGT AACTATGACTCTCTTAAGG TAGCC |
| target-022 | Cy5-labeled dsDNA cleavage for modified DNA assay | Forward template for Cy5-labeled modified dsDNA target amplification | GCCTACACGGGGAAGAGC GGGAGTTTAGTAACTCACC TTAAGGTAGCCGGTATTGC ATCG |
| target-024 | Cy5,FAM-labeled dsDNA cleavage | Cy5-labeled ssDNA for native dsDNA annealing | Cy5-GACGAGGCATTTGGCTACC |

| | | | |
|---|---|---|---|
| | | | TTAAGAGAGTCATAGTTAC TCCCGCCGTTTACCCGCG |
| target-025 | Cy5,FAM-labeled dsDNA cleavage | FAM-labeled ssDNA for native dsDNA annealing | FAM-CAAGCGCGGGTAAACGGC GGGAGTAACTATGACTCTC TTAAGGTAGCCAAATGCCT CGTC |
| target-026 | Cy5-labeled dsDNA cleavage for bubble DNA assay | Cy5-labeled ssDNA for Bubble1 dsDNA annealing | Cy5-GACGAGGCATTTGGCTACC CCCAGAGAGTCATAGTTAC TCCCGCCGTTTACCCGCGC TTG |
| target-027 | Cy5-labeled dsDNA cleavage for bubble DNA assay | Non-labeled ssDNA for Bubble1 dsDNA annealing | CAAGCGCGGGTAAACGGC GGGAGTAACTATGACTCTC TCCCGGTAGCCAAATGCCT CGTC |
| target-028 | Cy5-labeled dsDNA cleavage for bubble DNA assay | Cy5-labeled ssDNA for Bubble2 dsDNA annealing | CAAGCGCGGGTAAACGGC GGGAGTAACTATGACTCTC TATTGGTAGCCAAATGCCT CGTC |
| Target-029 | Cy5-labeled dsDNA cleavage for bubble DNA assay | Non-labeled ssDNA for GC-rich dsDNA annealing | CAAGCGCGGGTAAACGGC GGGAGTAACTATGACTCTC TGGGGGTAGCCAAATGCCT CGTC |

**Table S4. The top 100 reads ranked by fold-change in deep sequencing.**

| Window 2 (7-12 nt) | Window 3 (13-18 nt) | Window 4 (19-24 nt) | Window 5 (25-30 nt) | Window 6 (31-36 nt) | Window 7 (37-42 nt) | Window 8 (43-48 nt) |
|---|---|---|---|---|---|---|
| TATGCC | ACGTGC | GTCGTT | CATCGC | GCTCAT | CCTAAT | GATAAG |
| TCTGAT | GACTGC | GGTACG | GGTCTC | CTGGAT | CCTAAC | ATAAAG |
| CGTACC | GCCTGC | GGAACA | AGAACT | TGGATG | TTTAAT | ATTAAG |
| AAGGCG | GTCAGC | CGCACC | TGCAAA | CTTCGG | CATAAT | ATTAAT |
| GGCGTC | GAAGGC | GCAGGT | TGAAGT | TTGTGA | GCTAAT | ATAAGA |
| GTCAGA | ACCCGC | TTATAT | TCTACT | GGCCGT | TCTAAT | GATAAT |
| GGTACA | TTTTGC | GTATCC | TCCGCT | GCCTCG | CCTAAG | ATTAAA |
| GCTAGC | ATCGGC | CGGAGA | GGGGAC | AGCATG | CTTAAT | ATTATA |
| GATGGA | GCATGC | AGTGAG | GCCGCC | ACAAGA | CCTAAA | GATATA |
| GAGTGC | TACGGC | TCCAGC | GACGCG | TTCTCG | GTTAAT | ATTAGA |
| CTGACT | TAAAGC | GTCGAA | CTCGCT | TCTCAT | CATAAC | ATAAGG |
| CAGTTC | TTATGC | GTACGC | CGCGTT | TCGTAG | TTTAAC | GCAGGA |
| CAGTGT | TAATGC | GGCACA | CAAGCC | GTAGAC | CTTAAC | GATAAA |
| ATCGAC | ACATGC | GCTTCC | ATGGCA | GATACA | CCTACA | GATAGA |
| AGGCCC | AACTAC | ATGTGA | ATGCTC | TGGTAG | GATAAT | ATTAGG |
| AGCATG | TACTGC | AGAGGC | AGTCGT | TGGCTA | GTTAAC | ATTATT |
| ACGTCG | ATAGGC | ACGTGT | AACAGT | TCGTCG | ATTAAT | GATACA |
| AAAGGC | TATTGC | GCGAGT | ACGCCT | GTTTGC | GCTAAC | GATATG |
| TTGAAT | AATAGC | GGCGCT | TTTGTC | GTTCGC | CCAAAT | ATTAAC |
| TGGCGT | ATCTGC | ACCGTG | TGAGAC | GTTCGA | TATAAT | ATAATA |
| TGCGCC | GTTTGC | GTCACT | TCTCGG | GTTCCT | TCTAAC | GATATT |
| TCGGAT | GTATGC | GGGACA | TCGCCT | GGAGCG | CTTAAA | ATAAAT |
| TCAGTA | TTAAGC | TGGATC | TCCGAT | GCTTCC | CTTACA | GTACTA |
| TATGAT | AATGGC | TGGACT | TCATTG | GCGCAT | CTCAAT | ATTATG |
| GAGCCG | TTCTGC | TCGTCC | TAGTGT | GAATCG | CCCAAT | ATTACA |
| CTGGTA | TAGAGC | GCGAGA | GTTGAC | GAAGGC | ATTAAC | GATAGG |
| CGGATG | ATGTGC | GCCATC | GTTCAT | CGTTAC | CATAAG | GTCCCC |
| CGGAGT | ATATGC | GATACA | GGCGAC | CAACAC | CTGAAT | ATAACA |
| CGAACT | GATTGC | CGAACG | GCCTGT | AGCCGA | GATACA | GATAAC |
| CCTGGA | ACAAGC | CAGCGT | GAGCCT | ACTCCG | TTTAAG | GATTAA |
| CCGTTG | ACCTGC | CAGCGA | GAGACT | TTGGAG | ACTAAT | GTCACA |
| CCGACC | ACTTGC | AGGAGA | GACGCT | GCTGGA | CGTAAT | ATTACG |
| CCAGTG | AACAGC | AGCCGA | GAAGGA | CAGGTT | CTAAAC | ATTAGT |
| CATGTC | GAATGC | ACAAGT | CGGGGA | CAGCTG | GTTAAG | GATTAG |
| ATTGTG | GAGTGC | AATACG | CCGTAT | AGTTGC | TTAAAT | ATAAAC |
| ATTACT | ATTGGC | AACACT | CCCGTA | ACGCTG | TATAAC | ATAATG |
| ACCGAT | ACTAGC | GGAACT | CCATGC | TGCCGC | CTAAAT | GATCGC |

| | | | | | | |
|---|---|---|---|---|---|---|
| TCGTCT | GTCTGC | GGGACT | CAGGGA | ATTTTA | CTCAAC | GCAAAG |
| GGGACC | ATAAGC | GGGTGT | CACCTA | GCGTAC | CTGAAC | GATATC |
| CGTCTT | AGGTGC | GGCTCT | ATTTCT | CTGTGG | GTTAAA | ATAACG |
| CGGACC | TTTCGC | GGATCT | ATTGTC | TTCGTT | CCAAAC | ATAAGT |
| CATGTA | GTTCGC | GGGGGT | ATCCGG | TGTGTC | CCTTAT | GCTAAT |
| CGCTGC | ATTTGC | GGTACA | AGTCCG | TCGGCC | CCGAAT | GATAGT |
| AGAGGT | GTTAGC | GGCAGT | AGGTCT | TCGCGG | TTTACA | ATTTAA |
| TGCCGT | ATCAGC | GGAGGT | AGGTCG | GTGCCA | CACAAT | GATTAT |
| TTGACG | TTTGGC | GTGACC | AGGCGT | GTACCA | CGTAAC | GCTAAG |
| TTAGAC | GTTGGC | GGGTCT | AGGAGC | GGTTCA | GTAAAT | GTTAAG |
| TGGCGC | AATTGC | GGGAGA | AGCCGA | GGGATC | TTCAAT | GTTTAA |
| TGGCCT | AAGTGC | GTGACA | ACATTA | GCCCAT | CATAAA | GATACG |
| TGATTA | TCATGC | GGTACT | TGTCGT | GATTCC | GTTACA | GCTATA |
| TCGAGC | AAGAGC | GGTGGT | GGTCCT | GACTTA | CCATAT | ATTACT |
| TCCGTA | TAAGGC | GTTACT | TGTCCT | GACACT | CAAAAT | GGAGAA |
| TATGCG | CACAGC | GGAACC | TACGGT | CTCCCA | GCTAAG | ATAAGC |
| TAGTCC | AAAAGC | AGCACC | GTGCAG | CTACTC | TTGAAT | GCAAGG |
| TAGGCT | GTAGGC | GGCACC | AGGGTC | CGTATT | GATAAC | GCAATG |
| TACGGA | TGTTGC | GGTAGT | ACGTGA | CGCGAT | CTAAAG | GCTATG |
| GTCGCC | ATTCGC | GGATGT | TGGTCC | CCTTAG | TTTAAA | GCAGTA |
| GTATGA | TAGTGC | GTAACT | ACGTAG | CCCCGT | GTGAAC | ATTTAG |
| GGCTTC | TCTTGC | GCTATT | AAAGGC | CATGCG | CTCAAG | GCTATT |
| GGCGGC | GGTTGC | GTATCT | TGGCGC | CAGCCG | CCTTAC | ATATAA |
| GCGGTC | ACAGGC | GGTACC | GCATGA | CACCCA | CCTACT | ATTAGC |
| GCCTAC | TACAGC | GGAAGT | TTTGAC | CAAGTA | TCTAAG | GTATTA |
| GCCATG | AGTTGC | GGGTTT | TTGTAA | ATTTGT | TTAAAC | GCAAGA |
| GCCACT | ACTGGC | GTTACC | TTGCCC | ATCTAG | GTCAAT | ATAAAA |
| GCATAC | GACGGC | GTATGT | TTGATG | AGGGGA | GTGAAT | ATTGTA |
| GCAGCC | GTACGC | GGCGGT | TTGACC | AGGCGC | CATACA | ATATAG |
| GAGGCA | AGTGGC | GGGAGC | TTCTCT | ACGTGC | ATTAAG | GTTTAG |
| GAGCAG | CTTTGC | GTCAGT | TTCGTT | ACGGAC | ACTAAC | GATACT |
| GAGAAT | GTCGGC | GTCACC | TGTTCT | ACAGCA | CCTTAG | GTATAA |
| GACTTT | AGGAGC | GGGGCT | TGGGAA | CCGCAA | CCTCAT | GTAAAG |
| GACGGC | TATGGC | GTGTCT | TGGACG | TGCTTA | CTGAAG | ATTGAA |
| GACAGT | GTATAC | GGTATT | TGCACT | TGATAG | CTTACG | GATGTA |
| CTCCCT | TTAGGC | GTTAGT | TGACGA | TGACGC | GCTAAA | GCCATG |
| CGTGAC | GCTTGC | GTGGGT | TGACCT | TATCTG | CCCAAC | GTTTTA |
| CGTAGT | AGATGC | GGTGCT | TCTGTC | GTCGAC | CTAACA | ATAATT |
| CGGCTC | TTTAGC | GTGAGT | TCGGCT | GTATAG | GTAAAC | ATAGAG |
| CGATCG | GGGTGC | GGGATA | TCGCCA | GATGAC | CAACAT | ATTATC |

| | | | | | | |
|---|---|---|---|---|---|---|
| CGAGCC | GGATGC | CCCACC | TCCTTA | GATCGA | CTAAAA | GTTTTG |
| CCTTAG | GCTAGC | GGGACC | TCCACC | CGCAGT | CTCAAA | GCAATA |
| CAGGTA | GCTGGC | GGGGGA | TATCAC | ATTACG | CCATAC | ATATAT |
| CAGACA | TGAAGC | GGTAGA | TAGGTG | ATGCTC | GTCAAC | GCAACA |
| CACTAT | ATGCGC | GGTATA | TAGGCC | TGACTA | CAAAAC | GATTTA |
| CACGAT | GGAAGC | GGCTGT | TACACT | GCGGAG | CTTAAG | GTAATA |
| ATGTAC | AGTAGC | GGGTGC | TAACGT | CGCAGA | GGTAAT | GCTTTA |
| AGTTCT | TTCAGC | GGTATC | TAACAG | GCCGGC | CCATAG | GATGAG |
| AGTGCT | TATAGC | GGTTCT | GTTCGC | TGCATG | CGTAAG | ATAGAA |
| AGGCGA | CTATGC | GTAACC | GTTCAG | TTGTCA | CCGAAC | ATTGAT |
| AGGAGG | TAGTAC | GGGGTT | GTTCAC | TTATCG | TCTAAA | GCTTAT |
| AGCGCC | GTGAAC | GTAGGT | GTGTTA | TGTAGT | GTAAAG | GATTAC |
| AGATGC | GATCGC | GGGTGA | GTGGAC | TGCCTG | CACAAC | GTTAAA |
| AGAGGA | CCGTGC | GGTGTT | GTCGCC | TGCACA | TTAAAG | GTAATG |
| AGAACG | CCCAGC | GGCATT | GTCCAG | TGATTT | TTGAAC | ATTGTT |
| AGAAAT | CCAAGC | GTGAGA | GTAGCC | TGATTC | ATAAAT | GACCCG |
| ACTGGA | ATGTCC | GGCTTT | GGTATT | TGATAA | CTACAA | GATGAT |
| ACGGCA | ATACGC | GTAAGT | GGGTCC | TGACAG | AATAAT | ATTTAT |
| ACATCA | ACACGC | GTGACT | GGGACC | TCTGGA | TCAAAT | GTTATT |
| AATGTT | GATGGC | GGTAGC | GGCCTT | TCTGAG | CTGACA | ATTGAG |
| AATGCC | ACCGGC | GGTGGA | GGCAGA | TCGCGC | TTTACT | GATCAA |
| AAGTGC | TAACGC | GGAGGA | GGCAAC | TCGAAG | TGTAAT | GTTCTT |
| AAGTAC | ATCCGC | GGTTGT | GGATCT | TCCGTC | TTATAT | GTTATA |

**Table S5. Statistics for cryo-EM analysis and model refinement.**

| | 3'-RNA bound complex (pre-cleavage) | 3'-RNA bound complex (first-strand cleavage) | 5'-RNA and R2 complex | L-RNA bound complex |
|---|---|---|---|---|
| PDB ID | 8IBW | 8IBX | 8IBY | 8IBZ |
| EMDB ID | EMD-35347 | EMD-35348 | EMD-35349 | EMD-35350 |
| **Data collection and processing** | | | | |
| Microscope | Titan Krios | | | |
| Detector | Gatan K3 with GIF Quantum (20eV slit) | | | |
| CS (mm) | 0.01 | 0.01 | 0.01 | 2.7 |
| Magnification | 64K | 64K | 64K | 81K |
| Pixel size (Å) | 1.0979 | 1.0979 | 1.0979 | 1.0825 |
| Electron dost ($e^-$/ $Å^2$) | 50(32 frames) | 50(32 frames) | 50(32 frames) | 50(32 frames) |
| Defocus range (μm) | -1.5 ~ -2.0 | -1.5 ~ -2.0 | -1.5 ~ -2.0 | -1.5 ~ -2.0 |
| Micrograph Number | 5,039 | 5,039 | 4,502 | 4,647 |
| **Reconstruction** | | | | |
| Software | | | | |
| Particles picked | 3,557,515 | 3,557,515 | 3,544,913 | 2,299,896 |
| Particles refinement | 89,089 | 106,634 | 296,297 | 264,298 |
| Symmetry | C1 | C2 | C1 | C1 |
| Resolution (Å) | 3.60 | 3.74 | 3.47 | 3.04 |
| Sharpening B-factor ($Å^2$) | 136.5 | 158.6 | 162.0 | 109.2 |
| **Refinement** | | | | |
| Software | | | | |
| Model composition | | | | |
| Number of atoms | 8,913 | 8,690 | 9,660 | 9554 |
| Protein residues | 924 | 924 | 799 | 823 |
| Nucleotides | 124 | 113 | 201 | 189 |
| B factors (Protein/Nucleotide) | 80.73/167.24 | 104.02/176.51 | 72.97/132.46 | 46.60/40.14 |
| Bonds RMSD | | | | |
| Bonds lengths (Å) | 0.004 | 0.003 | 0.004 | 0.002 |
| Bonds angles (°) | 0.659 | 0.659 | 0.648 | 0.539 |
| **Validation** | | | | |
| MolProbity score | 1.98 | 2.01 | 1.98 | 1.57 |
| Clash score | 10.25 | 10.97 | 9.55 | 5.87 |
| Rotamer outliers (%) | 1.44 | 1.44 | 1.26 | 0.74 |
| C-beta outliers (%) | 0.00 | 0.00 | 0.00 | 0.00 |
| Ramachandran plot | | | | |
| Favored (%) | 95.31 | 95.29 | 94.09 | 96.33 |
| Allowed (%) | 4.69 | 4.71 | 5.91 | 3.67 |
| Outlier (%) | 0.00 | 0.00 | 0.00 | 0.00 |
| Model vs. Data | | | | |
| CC mask/box | 0.70/0.74 | 0.75/0.80 | 0.76/0.78 | 0.66/0.68 |

**Titles and legends for supplementary video**

**Video S1. Related to Figures 3-6.**

**3'-RNA and 5'-RNA bound complex.** EM maps and atomic models of 3'-RNA and 5'-RNA bound are sequentially presented. The conformation changes within the two conformations of 3'-RNA, and 5'-RNA bound complex are simulated in Chimera-X.

**Video S2. Related to Figure 6.**

**Simulation of sequential RNA binding.** Structures of L-RNA bound complex and 5'-RNA bound complex are used for these conformation changes simulated in Chimera-X.
